# An open structure-guided RNA foundation model for generalizable prediction

**DOI:** 10.1101/2025.08.06.668731

**Authors:** Heqin Zhu, Ruifeng Li, Ao Chang, Haobin Chen, Feng Zhang, Fenghe Tang, Tong Ye, Xin Li, Yunjie Gu, Peng Xiong, S. Kevin Zhou

## Abstract

RNA language models have achieved strong performance across diverse down-stream tasks by leveraging large-scale sequence data. However, RNA function is fundamentally shaped by hierarchical structure, making structural information important for pre-training. Existing approaches often depend on noisy annotations or task-specific objectives that can limit generalizability. Here we present structRFM, a structure-guided RNA foundation model pre-trained on 21 million RNA sequence–structure pairs by integrating base-pairing interactions into masked language modelling through a pair-matching operation. We also introduce MUSES, a multi-source ensemble of secondary structures designed to mitigate predictor bias, and a dynamic masking ratio that balances structure-guided and nucleotide-level masking. structRFM learns joint sequence and structure representations as sequence-level, nucleotide-level and pairwise features for broad downstream adaptation. It ranks among the leading models in zero-shot homology classification and establishes new benchmarks for secondary-structure prediction. Its tertiary-structure adaptation, Zfold, consistently improves both three-dimensional predictions and their extracted secondary structures, with performance comparable to AlphaFold3 [31] across CASP15-natural, CASP16 and RNA-Puzzles. structRFM also improves performance on function-prediction tasks and generalizes to newly released RNA families and long non-coding RNAs. Together, these results demonstrate the effectiveness of structure-guided pre-training and highlight multimodal RNA language models for robust structural and functional inference. The complete 21-million-pair dataset, model weights and source code are openly available to support multimodal RNA foundation-model development.

## 1 Introduction

Ribonucleic acid (RNA) not only serves as a carrier of genetic information from deoxyribonucleic acid (DNA) transcription to protein translation, but also folds into diverse structures and functions as a versatile regulator of cellular processes [5]. Beyond the classical roles of ribosomal RNA (rRNA) and transfer RNA (tRNA) in gene expression, numerous non-coding RNAs (ncRNAs) act as essential regulators, such as internal ribosome entry sites (IRESs) that mediate 5’ cap-independent translation in circular RNAs [6] and small nuclear RNAs that mediate pre-mRNA splicing [7]. These functions arise from their hierarchical structures where primary sequence motifs determine base-pairing patterns and tertiary folds, which in turn shape biochemical activities and interactions. Unravelling the relationships linking sequence to structure and ultimately to function is therefore crucial for understanding RNA biology and advancing RNA-centric therapeutics.

Decades of innovation in nucleic acid sequencing have produced an unprecedented volume of RNA sequence data [8], transforming our ability to map transcriptome complexity [9, 10]. Inspired by advances in natural language processing, RNA language models pre-trained in a task-agnostic, self-supervised manner can capture representative patterns and contextual relationships from large-scale sequential data. This capability supports applications in decoding RNA patterns, predicting tertiary structures [11, 12] and inferring functions [13, 14]. Most existing RNA language models are pre-trained on ncRNA sequences for general-purpose applications [15]. RNA-FM [1] and RiNALMo [2] employ standard BERT architectures, RNAErnie [16] incorporates motif patterns during pre-training, and AIDO.RNA [17] scales the model to 1.6 billion parameters. These models exhibit broad downstream generalizability. In contrast, specialized models focus on particular RNA types or objectives, including PlantRNA-FM [18] for plant RNA motifs, UTR-LM [13] for untranslated regions and mRNA functions, and SpliceBERT [14] for splice site prediction. Together, these models support diverse structural and functional tasks.

As RNA structure defines molecular properties, mediates biomolecular interactions and ultimately determines function [3, 4], incorporating structural information into pre-training offers substantial potential beyond sequence alone. Several recent models integrate encoded structural data. RNABERT [19] incorporates aligned Rfam seed structures into base embeddings, ERNIE-RNA [20] introduces a pairwise position matrix into the attention mechanism, and MP-RNA [21] and UTR-LM [13] use auxiliary losses to reconstruct masked secondary structures. Although these approaches represent important progress, they also have limitations. RNABERT depends on potentially inaccurate seed alignments, which may hinder family discrimination. ERNIE-RNA represents canonical interactions using simplified scores, for example 3 for GC, 2 for AU and 0.8 for GU pairs, which may not capture the complexity of RNA structural features. MP-RNA and UTR-LM rely on ViennaRNA-derived structures [22] and task-specific supervised objectives, diverging from task-agnostic self-supervision and potentially biasing representations towards secondary structure prediction. To address this gap, a pre-training strategy is needed that incorporates reliable structural information while retaining task agnosticism and broad downstream adaptability.

Motivated by this, we introduce **structRFM**, a fully open structure-guided RNA foundation model pre-trained using structure-guided masked language modelling (SgMLM). SgMLM implicitly encodes base-pairing interactions through informed masking patterns and jointly models sequential and structural dependencies. structRFM therefore harnesses structural priors while maintaining task-agnostic pretraining, enabling adaptation to zero-shot prediction, secondary and tertiary structure prediction, and multiple function-inference tasks (Fig. 1a). Our work makes three key contributions towards integrating structural knowledge into RNA foundation modelling.

**Fig. 1:**
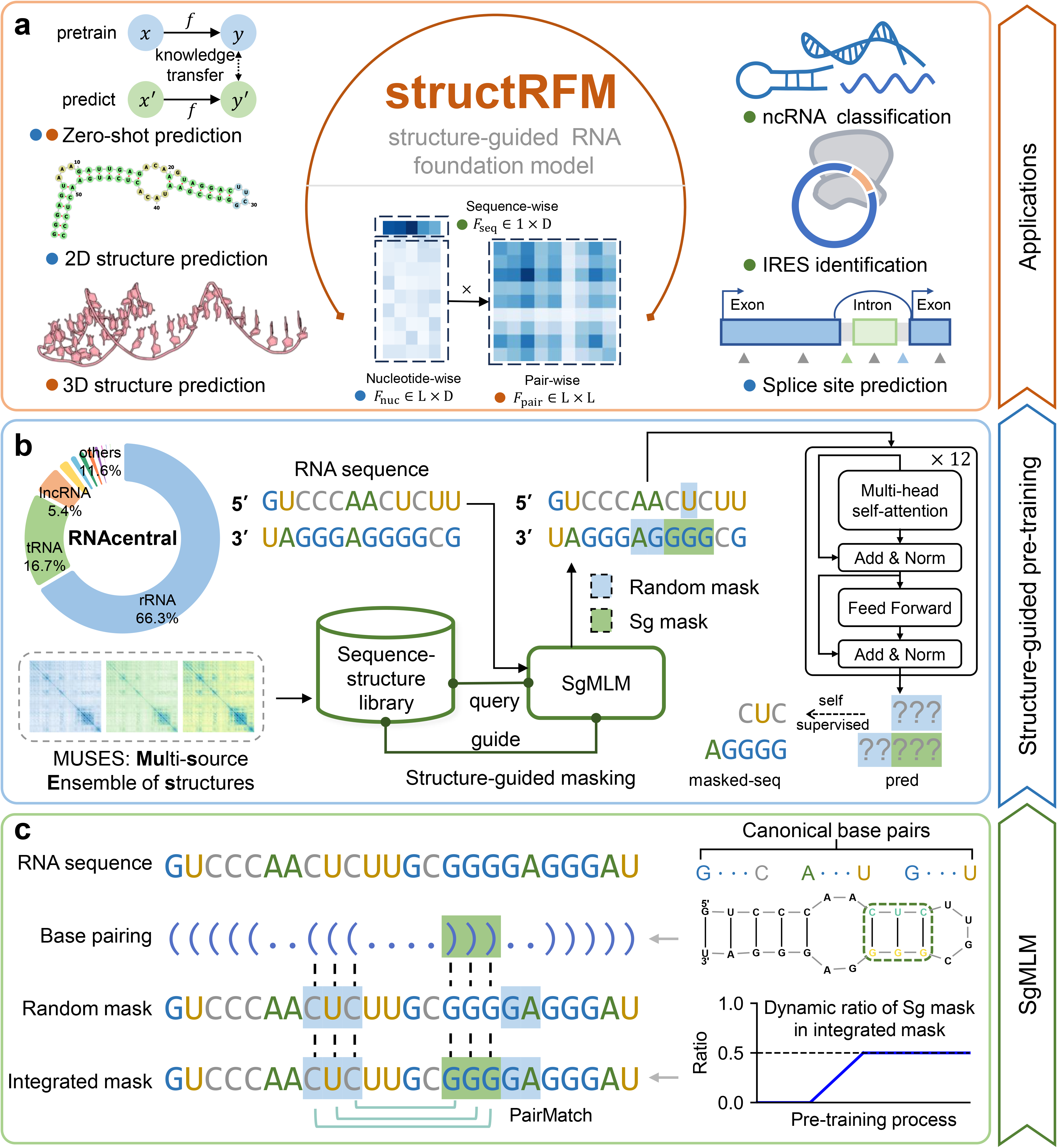
Overview of structRFM. **a** structRFM as an RNA foundation model for zero-shot or downstream-adapted structural and functional inference. The model outputs a sequence-level feature, a nucleotide-level feature and a pairwise feature, supporting tasks such as zero-shot homology classification, zero-shot 2D structure prediction, 2D and 3D structure prediction, splice site prediction, IRES identification and ncRNA classification. **b** Structure-guided pre-training workflow of structRFM. RNA sequences from RNAcentral are filtered by length, and their secondary structures are obtained using the proposed MUSES strategy, which uses a multi-source ensemble of secondary structures to construct a sequence–structure library for SgMLM pre-training. **c** The structure-guided masked language modelling (SgMLM) strategy. RNA base-pairing information is used to generate an integrated mask that combines random nucleotide-level and structure-guided masking, with the proportion of structure-guided masking dynamically increasing during pre-training.

First, we design SgMLM with two core components: structure-guided masking and dynamic masking balance (Fig. 1c). Structure-guided masking selectively masks tokens corresponding to canonical base pairs within local structural contexts, encouraging the model to recover paired nucleotides from surrounding loop regions without introducing a task-specific prediction objective. This implicitly guides structRFM to capture sequential patterns, base-pair relationships and hierarchical structural regularities, facilitating transfer to downstream structure and function tasks. The dynamic component gradually increases the proportion of structure-guided masking during pre-training, shifting the model from sequence-level learning towards a balance of nucleotide-wise and structure-wise masking. Furthermore, to address the complexity of RNA folding, we introduce MUSES (Multi-source ensemble of secondary structures), which integrates ViennaRNA [22, 23], CONTRAfold [24] and BPfold [25]. By combining thermodynamics-based, probabilistic and deep-learning-based predictors, MUSES mitigates biases arising from reliance on a single source and improves annotation robustness for pre-training.

Second, we comprehensively evaluate structRFM in zero-shot assessments and downstream-adapted structure and function tasks. structRFM ranks among the leading models in zero-shot homology classification and zero-shot secondary structure prediction within a comparison of 17 biological language models. It establishes new benchmarks for secondary structure prediction, while its tertiary-structure adaptation, termed Zfold, reduces *ɛ*RMSD and improves TM-score, GDT-TS and the INF of secondary structures extracted from predicted three-dimensional models across CASP15 [29], CASP16 [30] and RNA-Puzzles. These results support the effectiveness of SgMLM for structural adaptation. structRFM also transfers effectively to splice site prediction, IRES identification and ncRNA classification, achieving a top-k accuracy of 0.416 on SpliceAI [7] and F1 scores of 0.701 and 0.979 on the IRES [32, 33] and nRC [34] datasets, respectively. Furthermore, it generalizes to newly released RNA families, attaining INF values of approximately 0.65–0.70, and to long non-coding RNA classification with F1 scores above 0.94.

Third, we release the complete suite of resources, including the 21-million-pair sequence–structure dataset with ensemble base-pair annotations, the pre-trained structRFM model, fine-tuned models for structural and functional tasks, and source code. These resources are intended to support further development of multimodal biological foundation models. Overall, structRFM provides a fully open and extensible framework for robust downstream adaptation. SgMLM and the sequence–structure dataset are also extensible to other pre-training frameworks, empowering RNA foundation models with multimodal structural knowledge.

## 2 Results

In this section, we conduct a series of evaluations of structRFM, which was pre-trained on 21 million RNA sequence–structure pairs (Extended Data Fig. 6). We begin with feature visualization and zero-shot tasks to examine whether the learned embeddings encode family-level and base-pair information without labelled training data. We next evaluate secondary and tertiary structure prediction, including tertiary-structure quality and reliability. We then fine-tune structRFM on splice site prediction, IRES identification and ncRNA classification, spanning nucleotide- and sequence-level function prediction. Finally, we test generalization to newly released RNA families and unseen long non-coding RNA sequences from humans and mice. We denote the model pre-trained without secondary structures and SgMLM as structRFM*^−^*. Overall results and dataset summaries are provided in Extended Data Tables 1 and 2, respectively.

### 2.1 structRFM extracts category-discriminative and structure-encoded features

Task-agnostic self-supervised pre-training enables structRFM to capture intrinsic RNA sequence characteristics, while SgMLM enables the extraction of structure-encoded representations. For an input RNA sequence of length *L*, structRFM extracts a sequence-level feature *F*_seq_ *∈* ℝ^1^*^×D^* and nucleotide-wise features *F*_nuc_ *∈* ℝ*^L×D^*. Based on these representations, a pairwise feature *F*_pair_ *∈* ℝ*^L×L^* is obtained by averaging the 12 attention maps *F*_attn_ from the last transformer layer (*i* = 11, *j* = 0, 1*, …,* 11). This matrix encodes nucleotide-to-nucleotide relationships in a latent feature space.

To qualitatively assess the learned sequential and structural knowledge, we randomly sampled 23,994 RNA sequences across 19 RNA families from the pre-training dataset and extracted their sequence-level and pairwise features. We projected *F*_seq_ into two dimensions using t-SNE [35] and compared structRFM with a randomly initialized model without SgMLM pre-training. As shown in Fig. 2a, structRFM exhibited strong category-discriminative capability, separating different RNA families into distinct clusters, whereas the random model projected the same sequences into mixed representations in the latent space.

**Fig. 2:**
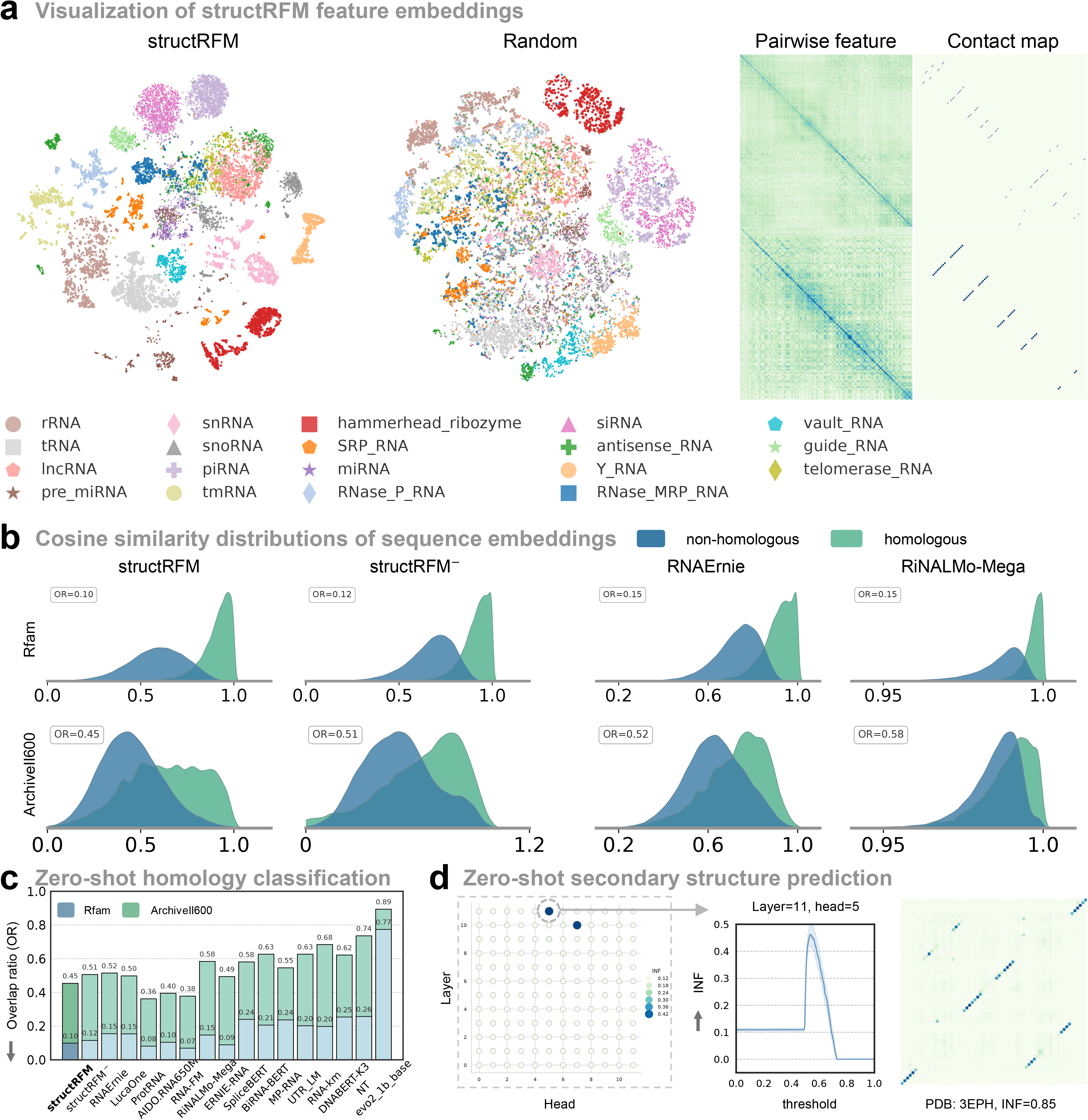
Feature visualization and zero-shot evaluation of structRFM. **a** Visualization of RNA embeddings from structRFM and a random baseline using t-SNE on a subset of RNAcentral (23,994 RNAs), showing that structRFM clearly separates RNA families. Also shown are two examples of pairwise features extracted from structRFM and their corresponding contact maps. **b** Cosine-similarity distributions between homologous and non-homologous RNA pairs for different models on the Rfam (24,523 RNAs) and ArchiveII600 (3,911 RNAs) datasets. **c** Overlap ratios (ORs) of the two distributions, showing the strong homology-discrimination ability of structRFM. The mean value is annotated for each method and dataset. **d** Assessment of structRFM for zero-shot secondary structure prediction. The first plot shows the INF metrics across attention maps from different layers and heads of structRFM, the second shows the INF metrics of the attention map from layer 11 and head 5 after post-processing across thresholds from 0 to 0.999 at intervals of 0.001, and the third shows an example attention heatmap. For the threshold scan, the solid line shows the mean INF across *n* = 116 independent RNA sequences, and the shaded band represents the 95% confidence interval.

Furthermore, we visualized *F*_pair_ as a heatmap and compared it with the native contact map. High-response regions aligned with base-pairing interactions, indicating that structRFM captures pairwise structural information. Collectively, these results show that structRFM learns representations encoding both family-level sequence properties and base-pair relationships, supporting downstream structure- and function-related tasks.

### 2.2 structRFM enables effective zero-shot prediction

Following previous work [15], we performed zero-shot homology classification and zero-shot secondary structure prediction to assess whether structRFM acquires generalizable structural and functional knowledge without task-specific labelled training data.

To evaluate whether sequence embeddings encode homology-level information, we randomly sampled 100,000 homologous and 100,000 non-homologous sequence pairs from each of the Rfam and ArchiveII600 datasets. In Rfam, homologous pairs belong to the same RNA family and non-homologous pairs belong to different families, whereas in ArchiveII600 homologous sequences share the same RNA type. We extracted sequence embeddings, computed the cosine similarity of each pair, and quantified the overlap ratio (OR) between the homologous and non-homologous distributions, where a lower OR indicates less overlap and better discrimination. As shown in Fig. 2b,c, structRFM ranked among the top four of 17 DNA, RNA and protein language models on both Rfam (OR = 0.10) and ArchiveII600 (OR = 0.45), showing clear separation despite AIDO.RNA having approximately tenfold more parameters. By contrast, DNA and specialized RNA language models performed less well, particularly on ArchiveII600 (OR *>* 0.6). These results suggest that general task-agnostic pre-training better captures family-level semantics. Additional similarity distributions are shown in Supplementary Figs. 1 and 2, and Extended Data Table 3 summarizes structRFM and other RNA language models.

Beyond homology classification, we assessed whether the model captures RNA secondary structure. Inspired by findings from RNA-MSM [36] that two-dimensional attention maps encode base-pairing patterns, we collected 116 RNA sequences from the PDB and extracted all 144 attention maps from the 12 layers and 12 heads of structRFM. Each attention map was treated as a base-to-base contact map and post-processed using thresholds from 0 to 1 at intervals of 0.001 before calculating INF. As shown in Fig. 2d, only a limited subset of layers and heads was informative for zero-shot secondary structure prediction, whereas other heads may primarily encode sequential information or joint sequence–structure relationships. Layer 11, head 5 achieved the highest INF of 0.462 at a threshold of 0.540, demonstrating a moderate ability to recover pairing features without task-specific supervision or labelled data. The sharp peak around the optimum further indicates that zero-shot contact prediction is sensitive to threshold selection when no calibration data are available.

### 2.3 structRFM achieves new benchmarks for secondary structure prediction

For secondary structure prediction, structRFM extracts the nucleotide-wise feature *F*_nuc_, which is concatenated with the one-hot encoding of the input RNA and processed by the MXfold2 structure module [37]. This module predicts four predefined stem– loop structural patterns, including helix stacking, unpaired regions, helix opening and helix closing. Zuker-style dynamic programming then converts these scores into the final secondary structure (Fig. 3a).

**Fig. 3:**
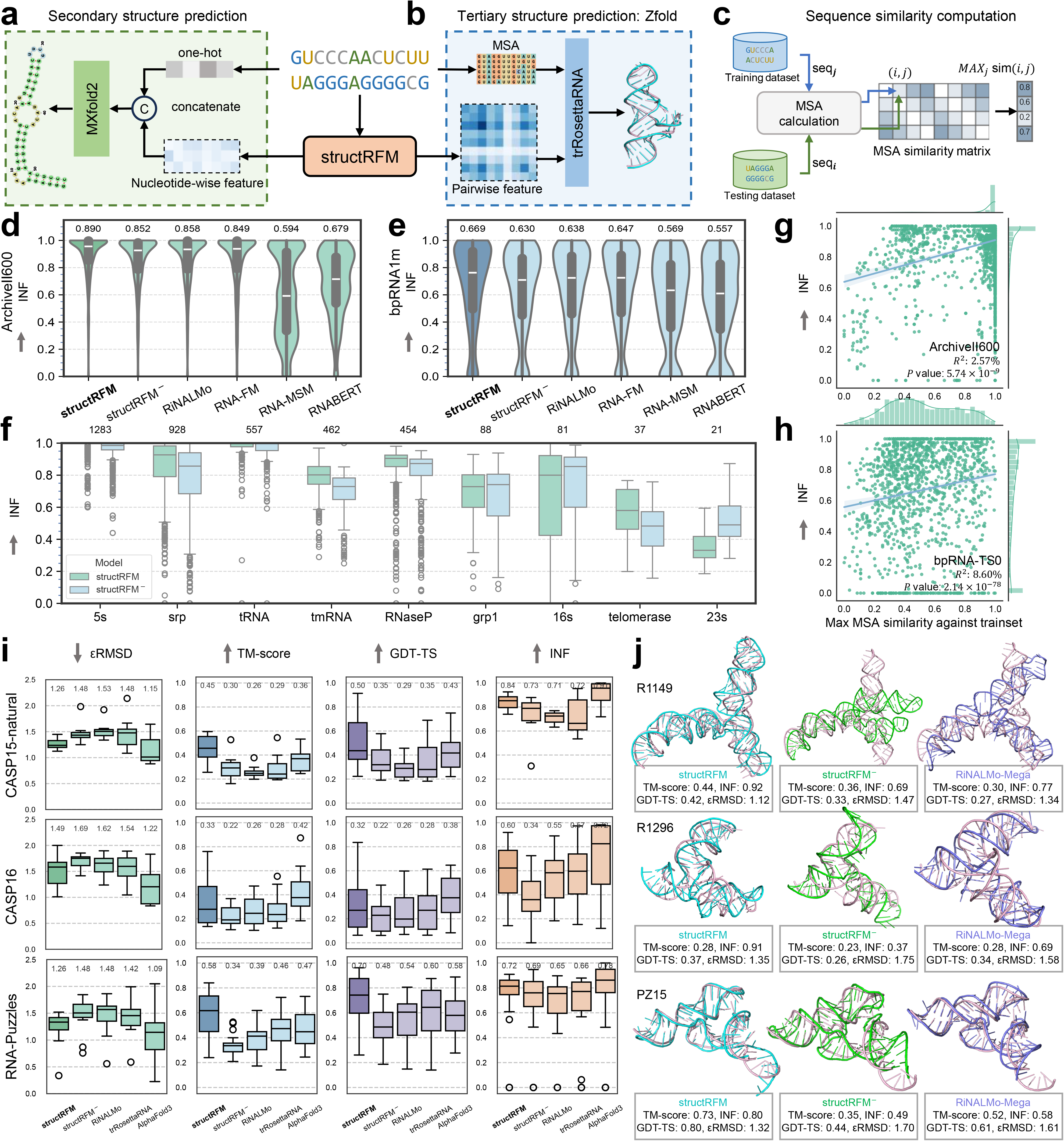
structRFM for structure prediction. **a, b** Neural-network architectures for secondary-and tertiary-structure prediction, respectively. **c** The pipeline for analysing sequence similarity between the test and training datasets. **d, e** INF comparisons of structRFM and other models for secondary-structure prediction on the ArchiveII600 (*n* = 3,911 independent RNA sequences) and bpRNA-TS0 datasets (*n* = 1,305 independent RNA sequences), respectively. In **d** and **e**, the inner box plots show the median (white centre line), 25th and 75th percentiles (lower and upper bounds of the thick line), and minimum and maximum (lower and upper thin whiskers); the mean is displayed above each violin. **f** INF comparison of structRFM and structRFM*^−^* on the nine families of ArchiveII600. The number of samples of each RNA family is displayed at the top of the figure. Centre lines indicate medians; lower and upper box bounds indicate the 25th and 75th percentiles; whiskers extend to the minimum and maximum observations within 1.5 times the interquartile range of the box bounds; and observations beyond the whiskers are plotted individually. **g, h** Relationships between INF and the sequence similarities of the ArchiveII600 and bpRNA-TS0 datasets to their corresponding training datasets, RNAStrAlign and bpRNA-TR0, respectively. Blue lines are linear regression fits, and shaded bands represent 95% confidence intervals. **i** *ɛ*RMSD, TM-score, GDT-TS and INF performance comparisons for tertiary-structure prediction on CASP15-natural (*n* = 8 independent RNA targets), CASP16 [30] (*n* = 13 independent RNA targets) and RNA-Puzzles (*n* = 20 independent RNA targets). Centre lines indicate medians; lower and upper box bounds indicate the 25th and 75th percentiles; whiskers extend to the most extreme observations within 1.5 times the interquartile range; observations beyond the whiskers are plotted individually; and values above the boxes indicate means. **j** Examples of predicted tertiary structures. Native structures are shown in light pink.

Following RNAErnie [16], we fine-tuned structRFM and baseline methods on RNAStrAlign [38] (19,313 RNAs) and bpRNA-1m [27] (12,114 RNAs), and evaluated them on ArchiveII600 [26] (3,911 RNAs) and bpRNA-TS0 [27] (1,305 RNAs), respectively. All models were evaluated under the same configuration for a fair comparison. As shown in Fig. 3d,e and Supplementary Table 1, structRFM achieved the highest performance among fine-tuned RNA language models, specialized learning-based methods and traditional methods. Specifically, structRFM reached INF values of 0.890 on ArchiveII600 and 0.669 on bpRNA-TS0, exceeding RiNALMo-Mega (0.858, *P* = 6.17 × 10^−15^, and 0.638, *P* = 0.0109, respectively; two-sided Welch’s t-test). It also outperformed task-specific and non-language-model baselines, including MXfold2 [37] (0.768 and 0.558) and ContextFold [39] (0.842 and 0.546). Compared with structRFM*^−^*, which lacks structure-guided pre-training, structRFM achieved INF values that were 4.46% higher on ArchiveII600 and 6.19% higher on bpRNA-TS0 (*P* = 2.74 × 10^−20^ and *P* = 1.10 × 10^−3^, respectively; two-sided Welch’s t-test). These results indicate that SgMLM imparts structural prior knowledge that facilitates downstream secondary structure prediction.

In addition, we evaluated family-level performance across the nine ArchiveII families (Fig. 3f and Supplementary Table 2). structRFM achieved higher INF across eight families with the largest sample sizes, indicating broad gains across RNA types. However, structRFM*^−^* slightly outperformed structRFM for 23S rRNA. This family has long sequences exceeding the 512-nt pre-training length, and existing secondary structure predictors also perform poorly as sequence length increases (Extended Data Fig. 1b,c).

Furthermore, we analysed sequence similarity between the training and test datasets. For each sequence, we used rMSA [40] and blastn [41] to generate multiple sequence alignments from RNAcentral [42] and NT [43], and defined test-to-training similarity as the maximum pairwise MSA similarity (Fig. 3c). ArchiveII600 showed higher training-set similarity than bpRNA-TS0. Nevertheless, similarity explained only 2.57% and 8.60% of the variation in INF, respectively (both *P <* 0.01), indicating weak linear relationships between sequence similarity and prediction accuracy (Fig. 3g,h).

### 2.4 structRFM enables robust and reliable tertiary structure prediction

RNA folds hierarchically, and tertiary structure is strongly constrained by secondary structure. We therefore hypothesized that accurate base-pairing information could improve tertiary prediction and that secondary structures extracted from predicted three-dimensional models could provide a complementary assessment of reliability. Moreover, unlike sparse secondary-structure representations such as dot-bracket notation or binary contact maps, language-model embeddings jointly encode sequence context and pairwise structural information and may therefore better support tertiary structure modelling.

As illustrated in Fig. 3b, we adopted trRosettaRNA [28] as the tertiary structure framework and used structRFM to extract a pairwise feature representing base-to-base relationships. The original trRosettaRNA pipeline combines multiple sequence alignments generated by rMSA [40] with a secondary-structure contact map predicted by SPOT-RNA [44, 45]. These inputs are used to predict one-dimensional orientations and two-dimensional contacts, distances and orientations, which are then converted into structural restraints for three-dimensional folding through energy minimization. In our variant, we replaced the predicted contact map with the pairwise feature from structRFM to assess whether richer structure-encoded embeddings improve tertiary prediction.

Following the original trRosettaRNA setup, we fine-tuned the resulting method, termed Zfold and referred to as structRFM for comparison, on 3,633 RNA chains from the PDB [46, 47] with a January 2022 cutoff and without self-distillation data. We evaluated it on CASP15 [29] (12 RNAs, including 8 natural and 4 synthetic targets), CASP16 [30] (13 RNAs) and RNA-Puzzles [48] (20 RNAs).

As shown in the target-level box plots in Fig. 3i and Supplementary Table 3, structRFM achieved mean *ɛ*RMSD values of 1.26, 1.49 and 1.26; TM-scores of 0.45, 0.33 and 0.58; GDT-TS values of 0.50, 0.32 and 0.70; and INF values of 0.84, 0.60 and 0.72 on CASP15-natural, CASP16 and RNA-Puzzles, respectively. It consistently outperformed RiNALMo-Mega, trRosettaRNA and structRFM*^−^* across the reported metrics, supporting the value of structure-guided pre-training for tertiary structure adaptation. Compared with AlphaFold3 [31], structRFM achieved slightly higher mean TM-score and GDT-TS on CASP15-natural and RNA-Puzzles. In the ablation, structRFM*^−^* was less structurally informative, yielding *ɛ*RMSD values of 1.48, 1.69 and 1.48 and INF values of 0.73, 0.34 and 0.69, respectively. For *ɛ*RMSD, the dataset-specific *P* values were 0.031, 0.057 and 0.012; for TM-score, they were 0.041, 0.074 and 1.90 × 10^−5^; and for INF, they were 0.081, 0.012 and 0.64 on CASP15, CASP16 and RNA-Puzzles, respectively. The corresponding target-level distributions and statistical comparisons are provided in Supplementary Table 3. RNA-Puzzles had higher average sequence similarity to the training data than CASP15 and CASP16, consistent with its stronger overall TM-score and INF (Fig. 4a). Nevertheless, for approximately one third of CASP15 and CASP16 targets with sequence similarity below 0.4, structRFM still achieved INF above 0.7 and TM-score above 0.4, demonstrating generalization when homologues or structural templates were limited. structRFM also outperformed structRFM*^−^* across the datasets and reported metrics, with particularly clear advantages at low similarity. Relative to trRosettaRNA, it increased secondary-structure INF by 16.7% on CASP15-natural, 5.26% on CASP16 and 10.8% on RNA-Puzzles, indicating benefits in difficult or low-homology settings. Representative examples R1149 from CASP15, R1296 from CASP16 and PZ15 from RNA-Puzzles further illustrate the improved tertiary structures and extracted base-pair patterns (Fig. 3j).

**Fig. 4:**
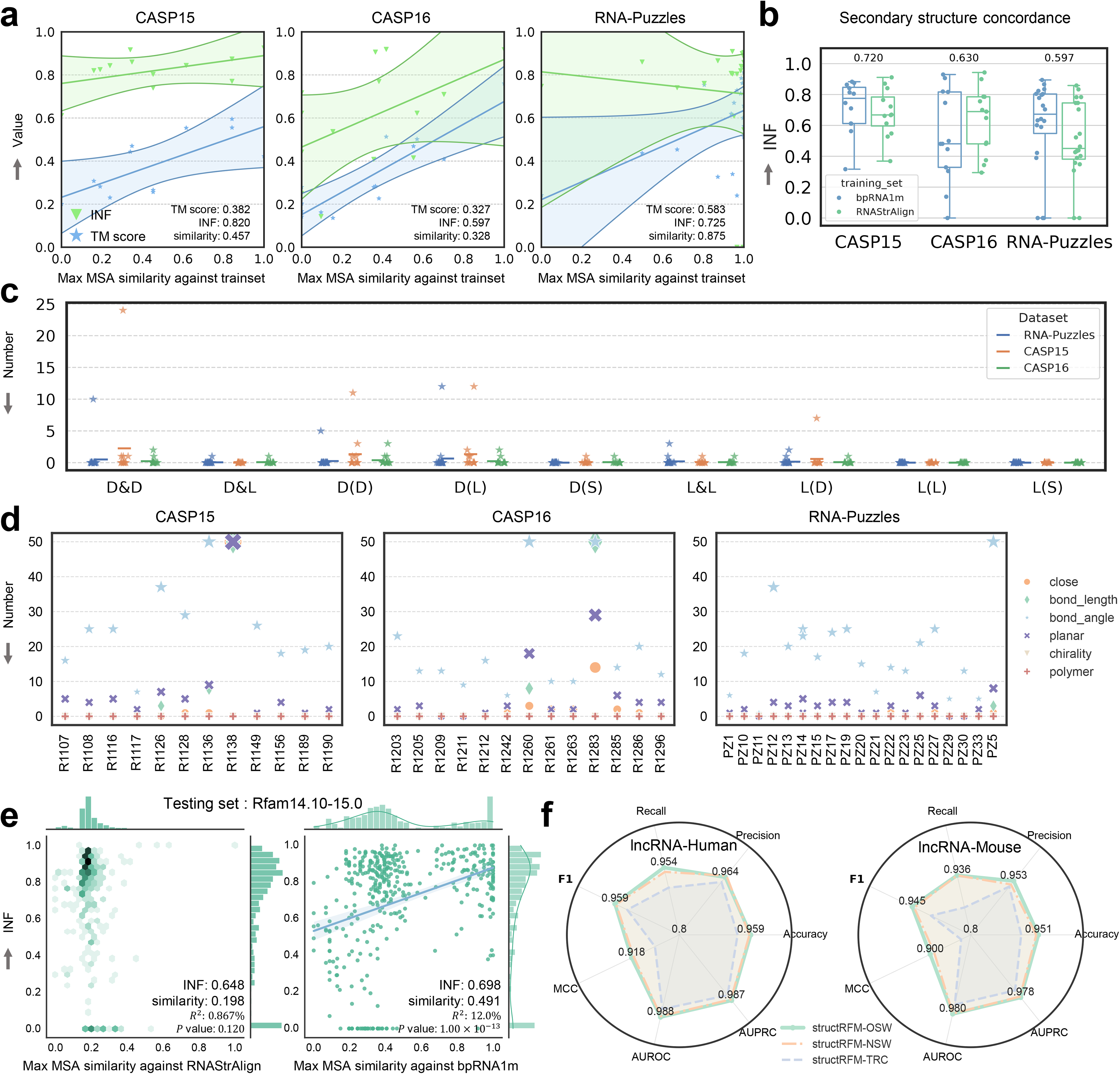
Analysis of predicted structures from the CASP15, CASP16 and RNA-Puzzles datasets and generalizability of structRFM. **a** Relationships of INF and TM-score with the sequence similarities of CASP15 (*n* = 12 independent RNA targets), CASP16 (*n* = 13 independent RNA targets) and RNA-Puzzles (*n* = 20 independent RNA targets) to the PDB training dataset. Centre lines are linear regression fits, and the surrounding shaded bands and boundary curves represent 99% confidence intervals. **b** Concordance between the secondary structures predicted by models fine-tuned for secondary-structure prediction (on RNAStrAlign and bpRNA-1m, respectively) and those reconstructed from the tertiary structures predicted by structRFM for the three datasets. For each comparison in **b**, *n* = 12 independent RNA targets for CASP15, *n* = 13 independent RNA targets for CASP16, and *n* = 20 independent RNA targets for RNA-Puzzles. Centre lines indicate medians; lower and upper box bounds indicate the 25th and 75th percentiles; whiskers extend from the minimum to the maximum; and points show individual RNA targets. **c** Entanglement analysis of structures predicted by structRFM on the three datasets. **d** Stereochemical quality analysis of the structures predicted by structRFM on the three datasets. **e** Generalization to new RNA families from Rfam v15.0. For comparisons against RNAStrAlign and bpRNA-1m, respectively, *n* = 280 and *n* = 436 independent RNA sequences with MSA results. Blue lines are linear regression fits, and shaded bands represent 95% confidence intervals. Two-sided Pearson correlation tests gave *r* = 0.093, *R*^2^ = 0.00867, *df* = 278, *P* = 0.120 and *r* = 0.346, *R*^2^ = 0.120, *df* = 434, *P* = 1.00×10^−13^, respectively. No adjustment for multiple comparisons was applied. **f** Evaluation of structRFM generalization to long non-coding RNA sequences from humans (21,605 RNAs) and mice (8,921 RNAs).

To further validate these results, we performed target-wise head-to-head comparisons of structRFM with four other methods, which showed consistent advantages across the three datasets (Extended Data Fig. 2a). We also evaluated aligned all-atom RMSD. structRFM achieved RMSDs of 9.97 Å on CASP15-natural and 5.32 Å on RNA-Puzzles, compared with 15.7 Å and 7.89 Å for AlphaFold3, respectively; its RMSD on CASP16 was 28.6 Å (Extended Data Fig. 2b). Although RMSD becomes difficult to interpret at large values and is affected by sequence length, the overall comparisons were consistent with the other structure metrics.

To provide a case-level assessment, we visualized all three-dimensional structures predicted for CASP15 and their extracted secondary structures together with the native structures (Extended Data Fig. 3). structRFM achieved a mean *ɛ*RMSD of 1.26 for the eight natural targets, compared with 1.41 for the four synthetic targets. The lower performance on synthetic RNAs may reflect their longer sequences and denser long-range interactions, which are underrepresented in the training distribution. Although overall predictions were strong, the extracted secondary structures sometimes missed short local stems and formed larger incorrect multiloops. CASP16 visualizations showed similar difficulties and reduced performance as sequence length increased (Extended Data Fig. 4), highlighting the remaining challenge of long-sequence modelling.

In addition, we assessed concordance between secondary structures predicted by models fine-tuned for secondary structure prediction and those extracted from tertiary predictions. Across CASP15, CASP16 and RNA-Puzzles, the former achieved average INF values of 0.720, 0.630 and 0.597, respectively, whereas the extracted structures achieved 0.822, 0.597 and 0.725 (Fig. 4b), indicating the generalizability and transferability of structRFM. The limited differences between models fine-tuned on RNAStrAlign and bpRNA-1m indicate that the choice of secondary-structure fine-tuning dataset had little effect on this comparison.

Finally, we evaluated the geometric, topological and stereochemical properties of the predicted tertiary structures [49–51]. Nearly all targets showed zero entanglement across most of the nine assessed categories, with CASP16 predictions containing fewer entanglements than CASP15 and RNA-Puzzles; no knotted artefacts were detected. However, bond-angle and planarity errors were common, whereas polymer and chirality errors were absent (Fig. 4c,d). These observations were consistent with the visualized tertiary and extracted secondary structures (Extended Data Fig. 3 and 4). They suggest that deep-learning-predicted structures may benefit from post-processing with physical force fields. Detailed category-level distributions and the analysis of the difficult CASP15 target R1138 are provided in Supplementary Information.

In summary, structure-guided pre-training consistently improves both secondary and tertiary structure prediction. Synthetic RNAs, long-range interactions and local stem formation remain challenging.

### 2.5 structRFM transfers across RNA function prediction tasks

We next evaluated whether structRFM transfers to three function-prediction tasks spanning nucleotide- and sequence-level outputs: splice site prediction, IRES identification and ncRNA classification.

At the nucleotide level, splice site prediction classifies each pre-mRNA position as a donor site, acceptor site or non-site. As shown in Fig. 5a, we used structRFM as the backbone and added a linear layer that maps the concatenated input features to nucleotide-level splice site scores. We fine-tuned structRFM and structRFM*^−^* on the SpliceAI dataset [7], which contains 162,706 training and 16,505 test sequences of length 100, and followed the BEACON configuration [52]. Because these pre-mRNAs differ substantially from the ncRNAs used during pre-training, the encoder remained trainable to support effective knowledge transfer. structRFM achieved a top-k accuracy of 0.42, exceeding RNA-MSM (0.38), UTR-LM (0.37) and RNA-FM (0.35), while remaining below the task-specific SpliceBERT model (0.45) (Fig. 5d). Compared with structRFM*^−^*, structRFM improved donor and acceptor prediction accuracy by 1.09% and 13.5%, respectively, indicating transfer of structural and sequential knowledge from ncRNA pre-training to pre-mRNA. The averaged model responses further showed the relatively sparse and consistent occurrence of donor and acceptor sites relative to the dominant non-site background (Fig. 5g).

**Fig. 5:**
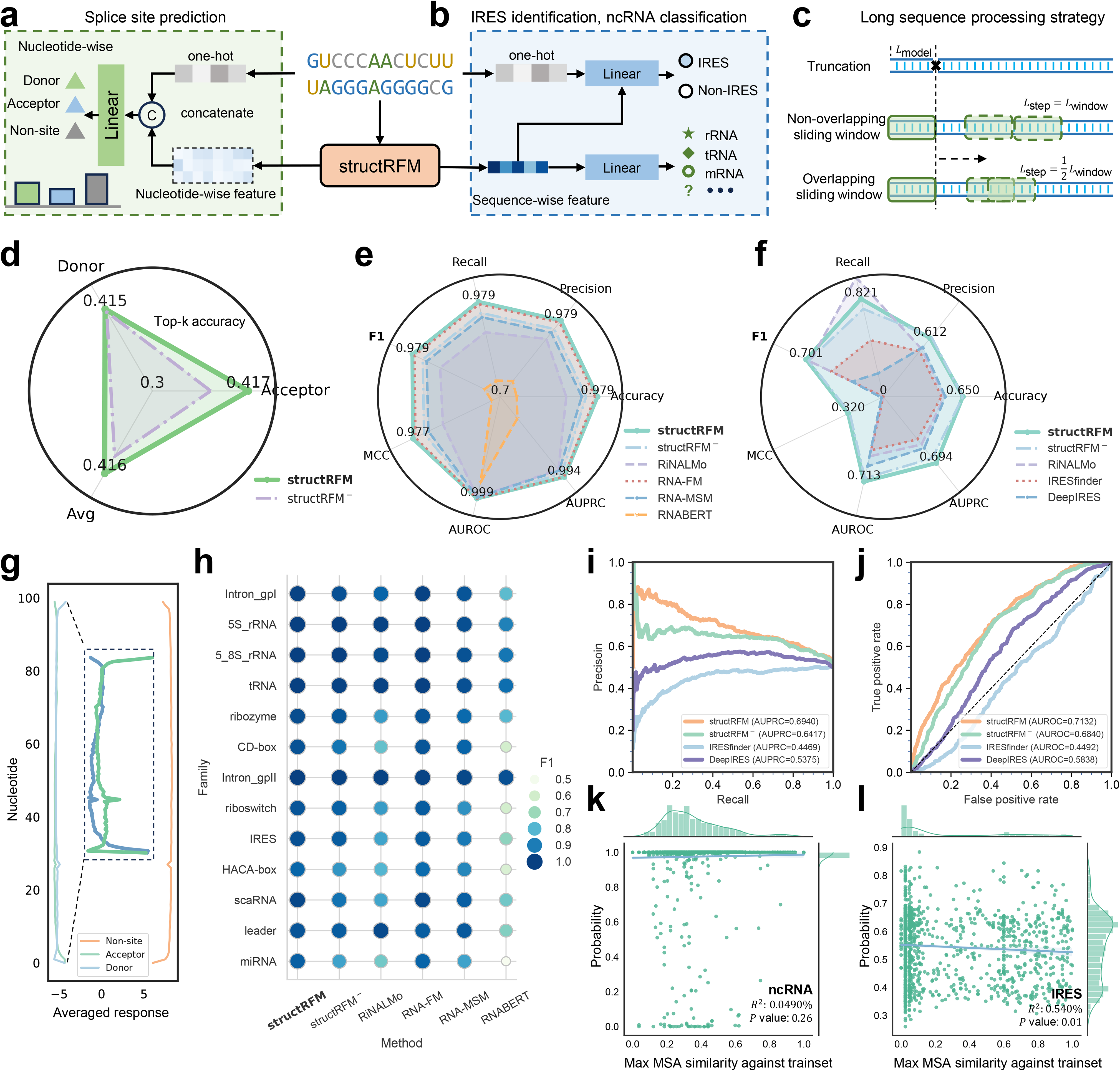
structRFM for function prediction. **a, b** Neural-network architectures for splice site prediction, IRES identification and RNA sequence classification. **c** Strategies for processing long RNA sequences, including truncation, non-overlapping sliding windows and overlapping sliding windows. **d** Top-k accuracy of structRFM and structRFM*^−^* for acceptor sites, donor sites and their average in splice site prediction (16,505 RNAs). **e, f** Radar plots of seven classification metrics for structRFM, the ablation model structRFM*^−^* and other methods for ncRNA classification on the nRC dataset (2,600 RNAs) and IRES identification (1,164 RNAs). **g** Prediction responses of structRFM across nucleotides for splice site prediction. Lines show mean predicted probabilities across *n* = 16,505 independent RNA sequences, and shaded bands represent 95% confidence intervals. **h** F1 comparison of structRFM with other models across the 13 ncRNA classes in the nRC dataset. structRFM and RNA-FM achieve state-of-the-art performance and outperform the other models by a substantial margin. **i, j** PR and ROC curves for IRES identification (1,164 RNAs). **k, l** Maximum MSA similarities of the ncRNA (*n* = 2,599 independent RNA sequences with MSA results) and IRES (*n* = 1,164 independent RNA sequences) test datasets to their corresponding training datasets, respectively, and the relationships between prediction confidence (probability) and sequence similarity. Blue lines are linear regression fits, and shaded bands represent 95% confidence intervals.

For sequence-level functional inference, IRESs mediate 5’ cap-independent translation in circular RNAs and can form stable, low-free-energy structures with potential applications in RNA therapeutics. As illustrated in Fig. 5b, we used the structRFM sequence-level feature with a feed-forward classifier. To address class imbalance, we trained models using complementary oversampling and undersampling strategies and combined their predictions by soft voting. We evaluated structRFM, its ablation and two specialized methods on an IRES dataset [32, 33] containing 12,998 training sequences and 1,164 balanced test sequences. structRFM achieved an AUPRC of 0.694 and an AUROC of 0.713, compared with 0.642 and 0.684 for structRFM*^−^*, respectively (Fig. 5i,j and Supplementary Table 6), supporting the contribution of SgMLM to capturing structural patterns relevant to IRES function. Furthermore, we compared F1 score, recall, precision, accuracy, AUPRC, AUROC and Matthews correlation coefficient across methods (Fig. 5f and Extended Data Fig. 5a). structRFM generally outperformed IRESfinder [53] and DeepIRES [54] across these metrics. It achieved an F1 score of 0.701, which was 47.5% higher than IRESfinder (0.475) and 146% higher than DeepIRES (0.284). Sequence similarity to the training set explained little variation in prediction confidence (*R*^2^ = 0.0540%, *P* = 0.01; Fig. 5l), indicating that the performance was not primarily driven by close sequence matches.

Finally, motivated by the strong zero-shot homology results, we evaluated supervised ncRNA classification on the nRC dataset [34], which contains 13 primary classes. The model uses the structRFM sequence-level feature followed by a linear classifier, and all models were fine-tuned under the same configuration as RNAErnie. structRFM achieved an F1 score of 0.979, MCC of 0.977, AUROC of 0.999 and AUPRC of 0.994, outperforming structRFM*^−^* and establishing the strongest overall performance among the compared methods (Fig. 5e, Extended Data Fig. 5b and Supplementary Table 7). RNABERT and RNA-MSM obtained F1 scores of 0.750 and 0.934, respectively. RNA-FM achieved performance comparable to structRFM despite weaker results in secondary structure and splice site prediction, suggesting that it captures strong classification-relevant features. RNAErnie [16] and the task-specific ncRDense [55] achieved F1 scores of 0.964 and 0.951, respectively. Furthermore, across the 13 individual classes, structRFM remained accurate for rare categories such as miRNA, HACA-box and scaRNA, for which structRFM*^−^* performed less well (Fig. 5h). This contrast is notable because rRNA accounts for 66.3% of the pre-training dataset, whereas these families are much less represented. The class-wise PR and ROC curves further supported robust classification across RNA families (Extended Data Fig. 5c). Most test sequences had low similarity to the training set, and similarity showed no statistically significant linear relationship with prediction confidence (*R*^2^ = 0.490%, *P* = 0.26; Fig. 5k).

Together, these results demonstrate that structure-guided pre-training transfers across nucleotide-level and sequence-level function prediction, including pre-mRNA and circRNA tasks that differ from the ncRNA sequences used for pre-training.

### 2.6 structRFM generalizes to new RNA families and long sequences

Finally, we evaluated the generalizability of structRFM on novel RNA families and long sequences. Because pre-training used RNAcentral v24.0, released on 25 June 2024, we curated 436 sequences from newly released Rfam v15.0 families that were absent during pre-training. We evaluated secondary structure prediction using models finetuned on RNAStrAlign and bpRNA-1m and reported INF together with maximum MSA similarity to the corresponding fine-tuning set. As shown in Fig. 4e, structRFM achieved INF values of 0.648 and 0.698, respectively. These results were obtained despite low maximum MSA similarities to the fine-tuning data (0.198 and 0.491, respectively). For the bpRNA-1m comparison, sequence similarity explained 12.0% of the variation in INF. Collectively, these results show that structRFM retains strong secondary structure performance on temporally separated RNA families.

Modelling long RNA sequences remains challenging, with existing secondary structure predictors showing reduced accuracy as sequence length increases (Extended Data Fig. 1). We therefore evaluated lncRNA classification using human and mouse datasets [56]. Before length filtering, lncRNA-Human contained 86,419 training and 21,605 test RNAs, and lncRNA-Mouse contained 41,961 training and 10,491 test RNAs. Owing to computational constraints, we retained sequences up to 3,000 nt, yielding 72,768 training and 21,605 test RNAs for human and 29,625 training and 8,921 test RNAs for mouse.

As outlined in Fig. 5c, we compared three strategies for processing sequences beyond the 512-nt input limit. Truncation retains only the first 512 nt and is therefore restricted to sequence-level tasks. Non-overlapping windows divide the full sequence into 512-nt segments and can also support nucleotide-level outputs. Finally, 50%-overlapping 512-nt windows preserve additional context across segment boundaries. Both sliding-window strategies markedly outperformed truncation, and the overlapping strategy performed best, achieving F1 scores of 0.959 for human lncRNAs and 0.9455 for mouse lncRNAs (Fig. 4f). These results demonstrate that structRFM can be applied effectively to long sequence-classification tasks, although windowing does not directly resolve global long-range modelling.

## 3 Discussion

In this study, we introduced structRFM, a fully open RNA foundation model that integrates base-pairing information into task-agnostic masked language modelling. By jointly learning sequential and structural dependencies, structRFM produces discriminative sequence-level, nucleotide-level and pairwise features that support zero-shot inference and downstream structure and function prediction. Unlike approaches that introduce task-specific supervised objectives during pre-training, SgMLM incorporates structural priors through the masking process and therefore retains a general-purpose learning objective. Across the evaluated settings, it improved secondary and tertiary structure adaptation and transferred to splice site prediction, IRES identification and ncRNA classification. Evaluation on newly released RNA families and long non-coding RNAs further supported generalization beyond the pre-training distribution. The 21-million-pair sequence–structure dataset, pre-trained and fine-tuned models, and source code provide fully open resources that can also support structure-guided extensions of other biological language models.

Despite these achievements, several limitations remain. First, the overlapping sliding-window strategy enables classification of sequences longer than the 512-nt input limit but cannot capture global sequence relationships or long-range base interactions, potentially limiting structure-related applications. Pre-training and fine-tuning directly on longer sequences may address this limitation, although at substantial computational cost. Second, MUSES integrates three secondary structure predictors but currently focuses on canonical base pairs. Future extensions could incorporate non-canonical pairs, stacking interactions and motif-level structural knowledge to better represent RNA conformational diversity. Third, although SgMLM improves structure-encoded representations and downstream generalization, its pre-training efficiency and scalability have not been systematically evaluated. Fourth, because the structRFM secondary structure model outperforms the individual predictors used by MUSES (Supplementary Table 8), self-distillation could allow successive pre-training rounds to use improved structRFM-derived annotations.

Looking ahead, structRFM provides a basis for more explicit multimodal RNA models. Whereas the current model implicitly integrates base-pairing interactions within a single encoder, modality-specific encoders and contrastive sequence–structure pre-training could explicitly align RNA sequences with structural representations at scale. Such learning may improve transfer across structure, function and interaction tasks, particularly when labelled data are limited. Furthermore, efficient long-context architectures such as Mamba [58], which avoid the quadratic sequence-length scaling of transformers [57], may improve the modelling of long RNAs. Another emerging direction is to integrate diverse biomolecular modalities within unified foundation models. AlphaFold3 [31] extends structure prediction across proteins, nucleic acids and small molecules, whereas Evo 2 [10] and LucaOne [9] move towards unified molecular language modelling across biomolecular types. These efforts reflect a broader trend towards general-purpose models of biological systems. The convergence of structure-guided pre-training, contrastive learning and unified multimodal modelling may therefore support more transferable, interpretable and broadly applicable foundation models.

## 4 Methods

### 4.1 Overview of structRFM

In this work, we present structRFM (Fig. 1a), an RNA foundation model that is pre-trained on multimodal data consisting of non-coding RNA sequences together with their corresponding secondary structures for general-purpose downstream adaptation. We first construct the pre-training sequence–structure dataset by filtering RNA sequences from the RNAcentral dataset [42] and collecting an ensemble of secondary structures. Next, as illustrated in Fig. 1b, we design a structure-guided masked language modelling (SgMLM) strategy to integrate RNA secondary structures into RNA sequence modelling during task-agnostic pre-training. After structRFM is fully pretrained, it is capable of extracting representative structural features from an input RNA sequence; these features can further play a vital role in various downstream tasks, including structural and functional tasks, such as secondary structure prediction, tertiary structure prediction, non-coding RNA classification, IRES identification in circular RNAs, and splice site prediction.

### 4.2 Pre-training sequence–structure dataset construction

Starting with the largest non-coding RNA dataset, RNAcentral version 24.0 [42], and following existing RNA language models [1, 16], we filter 21,477,078 RNA sequences from the active part of the RNAcentral dataset with a maximum sequence length of 512. This differs from some methods [17, 59] that collect RNA sequences from both the active part and the inactive part, which is no longer maintained. After collecting the RNA sequences, we apply the MUSES strategy (described below) to generate an ensemble of RNA secondary structures using three different methodological paradigms. Using a fixed random seed of 42, we split these sequence–structure pairs into training and test datasets containing 20,403,223 and 1,073,855 samples, respectively. This multimodal dataset consists of a large number of RNA sequences and RNA secondary structures, computationally alleviating the shortage of structural data at the secondary-structure level and facilitating the development of multimodal, structure-enhanced RNA foundation models. Statistics of sequence length, RNA family numbers, base-pair interactions and confidence indices are visualized in Extended Data Fig. 6.

### 4.3 Multi-source ensemble of secondary structures (MUSES)

Relying on a single secondary structure predictor risks imprinting its biases onto the foundation model. RNA folding is inherently defined by rugged energy landscapes with numerous metastable states, strong dependence on environmental conditions, and frequent pseudoknots that are rarely captured by standard predictors. Representing each sequence by a sole deterministic structure therefore under-represents the full conformational diversity of RNA.

To address this limitation, we introduce an ensemble framework termed **MUSES** (*Multi-source ensemble of secondary structures*). Instead of relying on a single predictor, MUSES integrates hypotheses from diverse sources, including ViennaRNA RNAfold v2.0 [22, 23], CONTRAfold v2.02 [24] and BPfold v0.2.5 [25], thereby combining three distinct methodological paradigms. First, ViennaRNA RNAfold provides a thermodynamics-based prediction using its well-established minimum free-energy model. Second, CONTRAfold contributes a statistically learned perspective, leveraging conditional random fields trained on structural databases to identify probable motifs. Finally, BPfold introduces a deep-learning approach integrated with physical priors that is robust and accurate by means of a high-quality, full-coverage canonical base-pair motif library for predicting secondary structures with confidence indices that reveal potential interactions between canonical base pairs, including G-C, C-G, A-U, U-A, G-U and U-G. The consensus structures are derived through a weighted integration of these individual predictions, prioritizing base pairs predicted with high accuracy across multiple frameworks. This tripartite approach mitigates the inherent biases and limitations of any single method, ensuring a more accurate and reliable structural model for further applications.

Specifically, each method produces a pairwise secondary-structure matrix *M_i_* ∈ [0, 1]*^L×L^*, where *M_i_*(*j, k*) denotes the base-pair probability (∈ [0, 1]) or pairing status (∈ {0, 1}). The deterministic dot-bracket outputs of BPfold are converted into calibrated binary matrices, whereas probabilistic predictors such as ViennaRNA RNAfold and CONTRAfold directly yield base-pair probabilities. Formally, the ensemble base-pair probability matrix is defined as follows:

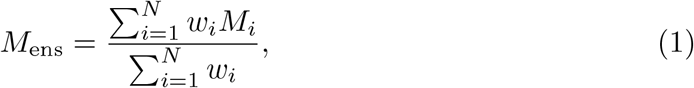

where *w_i_* reflects method-specific confidence (e.g., higher weight for pseudoknot-aware or thermodynamically consistent predictors). From *M*_ens_, a consensus secondary structure is derived using a maximum expected accuracy (MEA) decoder:

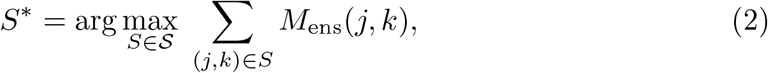

where is the set of valid secondary structures under RNA pairing constraints. This yields a consensus structure that reflects agreement across predictors while maintaining conformational diversity. Extended Data Fig. 1a presents our evaluation of MUSES across three datasets, with varying weights from 1 to 10 assigned to its three constituent methods. Ternary simplex plots illustrate performance variations across datasets and weight combinations, highlighting the limitations of relying on a single predictor. The final weights used to construct the pre-training dataset were set to 4:1:7 on the basis of the best MUSES performance across the three datasets.

### 4.4 Structure-guided masked language modelling strategy

To improve the ability of an RNA language model to learn base-pairing interactions and capture structural patterns, we carefully design a structure-guided masked language modelling strategy (SgMLM) to replace standard masked language modelling during the pre-training of structRFM, inspired by base-pair interactions. By doing so, we can leverage structure-level knowledge and extract discriminative representations of RNA sequences to support downstream tasks.

Specifically, given an RNA sequence **X** = *x*_1_*, x*_2_*, …, x_L_* ∈ {A, U, G, C, N}*^L^*, the standard masked language modelling strategy generates a random mask **M_r_** ∈ {0, 1}*^L^* for this sequence by sampling from a Bernoulli distribution with a probability of 15%, where **M_r_**(*i*) = 1 indicates that the token at position *i* is masked as “[MASK]”, and **M_r_**(*i*) = 0 indicates that the token at position *i* is left unchanged.

Based on this, we introduce the canonical base pairs of the RNA secondary structure into this procedure. The corresponding secondary structure of an RNA sequence with a length of *L* can be represented in the form of a binary adjacency matrix **C** ∈ {0, 1}*^L×L^*, where **C**(*i, j*) = 1 indicates that bases *i* and *j* form a base pair, and **C**(*i, j*) = 0 indicates that bases *i* and *j* do not form a base pair. As shown in Fig. 1b, we can obtain a secondary-structure mask **M_s_** ∈ {0, 1} *^L^* from the secondary structure, where **M_s_**(*i*) = 1 indicates that base *i* is part of a base pair and vice versa. The structure-guided masking strategy is designed to mask RNA bases with more paired interactions rather than isolated bases. Therefore, we apply an operation for matching canonical base pairs, denoted “PairMatch”, to account for these isolated bases in the random mask **M_r_** that participate in base-pairing interactions. The final structure-guided mask **M_f_** comprises two parts: (1) “PairMatch” applied to the “logical and” of the random mask **M_r_** and the secondary-structure mask **M_s_**; and (2) the “logical and” of the random mask **M_r_** and the “logical not” of the secondary-structure mask **M_s_**. This can be formulated as follows:

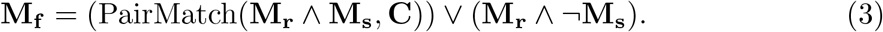

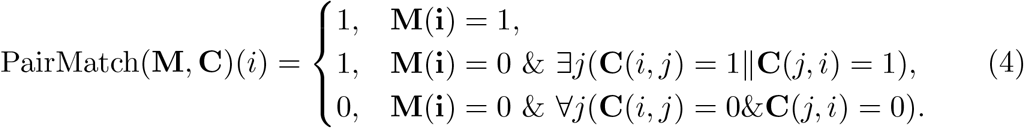

Throughout the training process, to conduct data augmentation and enhance the diversity of the training data, among the structure-guided masked tokens, we replace “[MASK]” tokens with random tokens 10% of the time and keep the masked input tokens unchanged for another 10% of the time. Furthermore, to enhance pre-training stability and enable structRFM to learn both token-wise and pairwise knowledge, we introduce a warm-up mechanism by gradually increasing the ratio *r* of structure-guided masking relative to standard masking from 0 to 0.5, which can be formulated as follows:

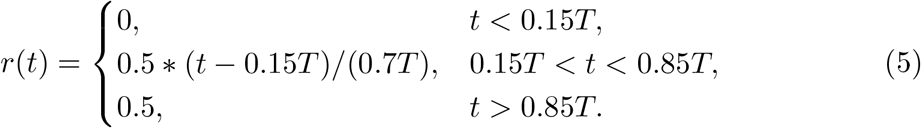

where *T* is the total number of training steps and *t* is the current training step. Therefore, the language model is ultimately pre-trained with both sequence-level and structure-level masking, jointly modelling the relationships between RNA sequences and base-pairing interactions and resulting in an enhanced capability to understand RNA structural patterns. During masked language modelling, the proxy task is to recover the masked tokens as their original nucleotide identities using a cross-entropy loss. By applying structure-guided masking, structRFM can infer masked matched base pairs from the surrounding unmasked, unpaired loop bases, intrinsically capturing base-pair interactions from the input RNA sequences, which is beneficial for structural prediction.

### 4.5 Pre-training structRFM with secondary structures

During the pre-training process, structRFM makes use of the constructed sequence– structure dataset. For an input RNA sequence, structRFM generates a structure-guided mask for it according to its secondary structure, tokenizes the input sequence, and embeds it as feature vectors. Consequently, structRFM utilizes a transformer [57] as its backbone, effectively models the long-range relationships in RNA sequences, and aims to learn the sequential and structural representations through the proxy task of recovering the masked tokens.

Specifically, for a given RNA sequence **X** = {*x*_1_*, x*_2_*, …, x_L_*} ∈ {A,U,G,C,N}*^L^* that is composed of the “A”, “U”, “G” and “C” nucleotides, with any other unknown nucleotide denoted as “N”, we replace the masked tokens with “[MASK]” tokens after applying structure-guided masking. Additionally, we add a “[CLS]” token and an “[SEP]” token at the beginning and end of the sequence, respectively, which is helpful for fine-tuning on classification tasks and for separating multiple input RNA sequences when dealing with RNA–RNA interactions and multiple chains. To keep the sequence length uniform within a batch, we pad it with “[PAD]” tokens at the end of the sequence. These tokens are further encoded as numbers and processed by an embedding layer to form the input sequence embedding *F*_0_ ∈ ℝ*^L×D^*.

As demonstrated in Fig. 1a, structRFM employs *N* consecutive transformer neural-network blocks as its backbone to process the input sequence embedding. The transformer block consists of a multi-head self-attention module (MHSA) and a feed-forward network (FFN), which is composed of linear layers and GELU activation. After the input feature is processed by the MHSA and FFN modules, the respective outputs are further processed using residual connections and layer normalization. The whole process can be formulated as follows:

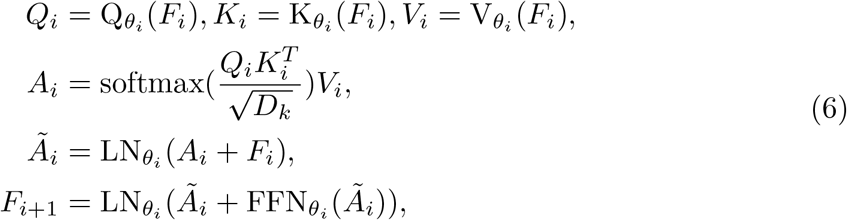

where *i* = 0, 1, 2*, …, N* 1 denotes the *i*-th transformer block with learnable parameters *θ_i_*.

### 4.6 structRFM feature embeddings for multitask inference

After pre-training, given an arbitrary RNA sequence of length *L* as input, struc-tRFM is capable of outputting a representative embedding *F* of shape ℝ^(1+^*^L^*^)^*^×D^* with encoded structural information. This embedding consists of a sequence-level feature and a nucleotide-wise feature. The sequence-level feature *F*_seq_ ∈ ℝ^1^*^×D^* is derived from the “[CLS]” token placed in the first row of the output embedding *F*, which is helpful for sequence-level classification tasks. The nucleotide-wise feature *F*_nuc_ ∈ ℝ*^L×D^* encodes each nucleotide of the RNA sequence and provides a powerful representation for nucleotide-level classification and regression tasks. Moreover, we derive a pairwise feature *F*_pair_ ∈ ℝ*^L×L^* by averaging the attention maps 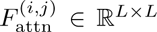 from the 12 attention heads in the last transformer layer (*i* = 11, *j* ∈ {0, 1*, …,* 11}). The resulting matrix encodes pairwise nucleotide-to-nucleotide relationships in a latent feature space and can be interpreted as a learned adjacency matrix, which is particularly important for modelling interactions between nucleotide pairs in structure prediction tasks. Formally, *F*_pair_ is computed as follows:

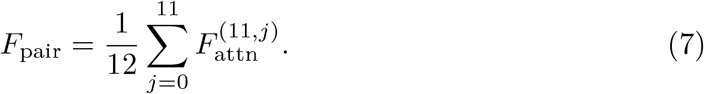

The above three kinds of feature embeddings, with discriminative structural information, are powerful representations that serve as transitional intermediates to be further processed by a simple multilayer perceptron or other downstream modules for various downstream RNA tasks, including structural and functional predictions with task-specific fine-tuning. For sequence-level classification tasks, such as non-coding RNA classification, the sequence-level feature *F*_seq_ is used. For sequence-level regression tasks, we use the mean of the nucleotide-wise feature *F*_nuc_ along the sequence-wise dimension. Besides, the nucleotide-wise feature *F*_nuc_ is intrinsically suitable for nucleotide-level tasks, such as splice site prediction. For RNA secondary-structure and tertiary-structure prediction, considering the difficulty of modelling interactions between base pairs, we employ the pairwise feature *F*_pair_ to provide sufficient base-to-base prior knowledge.

### 4.7 Fine-tuning structRFM for downstream adaptation

After pre-training structRFM with RNA sequences and their corresponding secondary structures in a task-agnostic, self-supervised manner, we are able to adapt it to various downstream tasks either through zero-shot prediction or by fine-tuning it with labelled data in a task-specific, supervised manner. Here, we demonstrate the supervised fine-tuning strategies in terms of fine-tuning different parts of task-specific neural networks and the ways of constructing neural architectures for various structural and functional tasks.

#### Fine-tuning strategy

The learnable parameters of structRFM for down-stream adaptation comprise two components: the pre-trained backbone (including the sequence-embedding layers and feature encoder) and task-specific heads or modules. These components give rise to two common fine-tuning strategies for transferring the knowledge learned by structRFM. The first strategy freezes the pre-trained backbone and fine-tunes the task-specific heads and modules, requiring fewer computational resources and less training time. The second strategy fine-tunes the entire neural network, including the backbone and task-specific components; it is more resource intensive but typically more effective. Given the distribution shift in the input RNA sequences, we used the second strategy in our experiments and compared the vanilla “structRFM” with its ablation variant “structRFM*^−^*”, which was pre-trained without base-pairing knowledge or the SgMLM strategy.

#### Neural architecture for downstream tasks

Given an input RNA sequence, structRFM, as a foundation model, processes it and extracts sequence-related and structure-encoded feature embeddings, which are further fed into the prediction heads to produce different output representations. Tasks such as RNA sequence classification only need a task-specific head—a fully connected layer—whereas some tasks that output complex dense predictions, such as secondary-structure prediction and tertiary-structure prediction, need post-processing procedures and structural modules.

##### Secondary structure prediction

RNA secondary structures exhibit base-pairing patterns and stem–loop regions, playing vital roles in modelling RNA tertiary structures and in the exploration of functional RNA motifs within non-coding genomic regions. Most existing methods predict the canonical base pairs (*i.e.*, A-U, G-C and G-U) of a given input RNA sequence of length *L*. The secondary structure can be represented as an adjacency matrix of shape *L × L*, in which each element stores the pairing relationship between base *i* and base *j*, *i.e.*, 0 for unpaired and 1 for paired. For predicting such dense connections in a scheme of nucleotide-wise classification, complex post-processing procedures, such as symmetry operations, sharp-loop removal and overlap removal, have been applied [60, 61], and thermodynamic priors have been adopted [25, 37] in previous methods. In our experiments, following RNAErnie [16], we adopt the lightweight MXfold2 neural network [37] as the output structural module. Specifically, as demonstrated in Fig. 3a, structRFM extracts the nucleotide-wise feature from the input RNA sequence and then concatenates it with the one-hot encoding of the RNA sequence, forming enhanced feature embeddings. The MXfold2 neural network further processes these feature embeddings and computes four kinds of folding scores at the base-pair level: helix stacking, unpaired region, helix opening and helix closing. Finally, a Zuker-style dynamic-programming approach [62] is applied during post-processing to predict the secondary structures.

##### Tertiary structure prediction

Regarding their widespread participation in cellular activities such as protein translation and gene regulation, RNA tertiary structures are essential for understanding their functions in biological processes. Predicting RNA tertiary structure remains challenging because of its dynamic nature and the lack of high-quality experimental data. An increasing number of methods [11, 12] use RNA language models to generate representative nucleotide-wise features as the input information for tertiary-structure prediction methods. In our experiments, as illustrated in Fig. 3b, we employ trRosettaRNA [28] as the structure module to replace the direct secondary-structure input with the pairwise feature produced by structRFM, aiming to examine the effectiveness of the pairwise feature generated by structRFM. The structural module takes multiple sequence alignments (MSAs) and the pairwise feature as inputs and outputs one- and two-dimensional geometries, which are further refined by PyRosetta [63] to generate the final tertiary structure.

##### Splice site prediction

Pre-mRNA splicing is an essential step in eukaryotic gene expression. Introns (non-coding regions) must be accurately removed, and exons (coding regions) are joined together by the spliceosome complex to form mature mRNA, which is then translated into a functional protein. Therefore, it is important to accurately detect the exon–intron boundaries—the splice sites. Splice site prediction is a nucleotide-wise classification task that aims to recognize whether each base is a donor site, an acceptor site or a non-site. As shown in Fig. 5a, we simply add a linear layer as the prediction head to the structRFM backbone to construct the neural-network architecture for splice site prediction. To compare with RNA language models benchmarked in BEACON [52], we follow the same configuration by passing the nucleotide-wise feature from the first layer of structRFM to a linear layer to make nucleotide-wise predictions.

##### IRES identification

The internal ribosome entry site (IRES), which mediates cap-independent translation in circular RNAs, is widely found in viral genomes and eukaryotic cells and exhibits high stability and tissue specificity. Therefore, accurate IRES identification aids in developing RNA therapeutics and targeted cancer interventions. As illustrated in Fig. 5b, the neural-network architecture for IRES identification is composed of the structRFM backbone and a simple classifier consisting of a fully connected linear layer. For this sequence-wise classification task, we adopt the sequence-wise feature extracted from the RNA sequence by structRFM and concatenate it with the one-hot encoding of the input sequence as the input feature for the classifier, making the architecture neat and effective. Furthermore, to deal with the imbalanced data, we fine-tune different models on an oversampled IRES dataset and an undersampled non-IRES dataset, respectively, and combine these models to make soft-voting predictions.

##### Sequence classification

Non-coding RNAs play various roles and demonstrate diverse functions in gene regulation and cellular processes. Accurate classification of ncRNAs is important for advancing biological research, developing RNA-based therapies, and improving disease diagnosis and treatment. In this downstream task, we simply use the sequence-wise feature provided by structRFM and a lightweight classifier head—a fully connected linear layer—as the neural-network architecture (Fig. 5b) to predict various ncRNA categories, such as rRNA, tRNA, mRNA and others.

### 4.8 Experimental configurations

At the pre-training stage, we pre-train structRFM for 1,627,466 steps with a batch size of 96 on two NVIDIA Tesla A100 80-GB graphics processing units for approximately 400 hours. structRFM adopts the vanilla BERT architecture [64], with a hidden dimension of 768 and the number of transformer layers being 12, and is pre-trained on our constructed sequence–structure dataset with a maximum sequence length of 512. This dataset consists of RNAcentral [42] sequences and secondary structures annotated using MUSES. We utilize the PyTorch [65] framework to implement structRFM and use the AdamW optimizer with a learning rate of 2×10^−4^. We design the SgMLM strategy and utilize the cross-entropy loss to enable structRFM to capture sequential and structural patterns.

We evaluate structRFM on two zero-shot assessments and five downstream finetuning tasks, namely zero-shot homology classification, zero-shot secondary structure prediction, secondary structure prediction, tertiary structure prediction, splice site prediction, IRES identification and ncRNA classification. The overall performance is presented in Extended Data Table 1. The experimental datasets, detailed in Supplementary Table 1, include Rfam [66, 67], PDB [46], ArchiveII600 and ArchiveII [26], bpRNA-1m [27], CASP15 [29], CASP16 [30], RNA-Puzzles [48, 68], Splice [7], IRES [32, 33] and nRC [34]. ArchiveII600 denotes the 3,911-sequence subset obtained by excluding the 64 sequences longer than 600 nt from the full ArchiveII dataset of 3,975 sequences; full ArchiveII was used only for the length-stratified method benchmark in Extended Data Fig. 1b,c.

For tertiary structure prediction, secondary structures are extracted from predicted tertiary structures using RNAView [69] and visualized using R2DT [70] and Forna [71]. We use nucleotide-interaction-centric *ɛ*RMSD [72] (lower is better), TM-score [73], and GDT-TS [74] to evaluate predicted tertiary structures, and Interaction Network Fidelity (INF, pair) [75] to evaluate the extracted secondary structures.

### 4.9 Statistics and reproducibility

All results were obtained from computational analyses of fixed benchmark datasets. No a priori power calculation was performed; sample sizes followed complete public benchmark partitions or published evaluation protocols, with all eligible records retained after the prespecified filters described in the Methods and Results. Each model-training and evaluation analysis was performed once under the reported configuration (one independent run); no independent experimental repeats or technical replicates were performed. Statistical summaries and hypothesis tests were calculated across distinct RNA sequences or structure targets, rather than across repeated model runs. Exact sample sizes and definitions of statistical displays are provided in the figure legends. Representative RNA structure visualizations were generated from the same single computational run and were not independently repeated. Blinding was not applicable because the study used fixed public computational datasets, predefined quantitative metrics and no experimental group allocation.

The custom structRFM, MUSES, SgMLM, PairMatch and Zfold code was version 0.0.8. Model and evaluation software comprised RNA LMs zero-shot evaluation v1 (https://zenodo.org/records/14430869), MXfold2 v0.1.0, trRosettaRNA v1.1, PyRosetta v2024.10+release.2c36cbc, rMSA code snapshot 2023-07-20, NCBI BLAST+ blastn v2.17.0+, RNAView v2.0.0, RNAErnie v1.0.0, RiNALMo-Mega code snapshot 2025-10-11, the AlphaFold3 web server accessed 2025-07-05, IRESfinder v1.1.0, DeepIRES code snapshot 2024-04-27 and SPOT-RNA code snapshot 2022-06-24. Analyses used Python v3.8.19, PyTorch v2.0.1, Transformers v4.45.1, NumPy v1.24.3, pandas v2.0.3, scikit-learn v1.3.0, SciPy v1.10.1, Matplotlib v3.7.2, seaborn v0.13.0, R2DT v2.0, PyMOL v2.5.0 and Forna v2015.12.02. Training and stochastic masking used a fixed random seed of 42.

## Data availability

The structRFM pre-training sequence–structure dataset is publicly available at Zenodo [76], while pre-trained model and fine-tuned model checkpoints are publicly available at GitHub (https://github.com/heqin-zhu/structRFM). The training and test datasets are publicly available at GitHub (https://github.com/heqin-zhu/structRFM), which are from the following sources: RNA-central v24.0 [42]: https://ftp.ebi.ac.uk/pub/databases/RNAcentral/releases/24.0/; Rfam v14.10 [66, 67]: https://ftp.ebi.ac.uk/pub/databases/Rfam/14.10/ and Rfam v15.0: https://ftp.ebi.ac.uk/pub/databases/Rfam/15.0/; ArchiveII600 and ArchiveII [26]: https://rna.urmc.rochester.edu/publications.html; RNAStrAlign [38]: https://github.com/mxfold/mxfold2/releases/tag/v0.1.0; bpRNA-1m, bpRNA-TR0 and bpRNA-TS0 [27]: https://bprna.cgrb.oregonstate.edu/download.php#bpRNA; PDB [46, 47]: https://www.rcsb.org/; CASP15 [29]: https://predictioncenter.org/download_area/CASP15; CASP16 [30]: https://predictioncenter.org/download_area/CASP16; RNA-Puzzles [48]: https://www.rnapuzzles.org/puzzles/list/; SpliceAI [7]: https://github.com/Illumina/SpliceAI; IRES [32, 33]: GEO GSE74277 (https://www.ncbi.nlm.nih.gov/geo/query/acc.cgi?acc=GSE74277) and GSE178718 (https://www.ncbi.nlm.nih.gov/geo/query/acc.cgi?acc=GSE178718); nRC [34]: http://tblab.pa.icar.cnr.it/public/nRC/paper_dataset/; lncRNA-Human (GEN-CODE 2021) [56]: https://www.gencodegenes.org/human/; lncRNA-Mouse (GENCODE 2021) [56]: https://www.gencodegenes.org/mouse/; and the NCBI nucleotide (nt) database [43]: https://ftp.ncbi.nlm.nih.gov/blast/db/. The study-specific processed IRES split and source data underlying the figures are provided with this paper. All other data are available within the article and its Supplementary Information. No restrictions apply.

## Code availability

The source code of structRFM is publicly available at Zenodo [77] and GitHub (https://github.com/heqin-zhu/structRFM).

## Acknowledgements

We thank Prof. Zaixi Zhang (Assistant professor, The Hong Kong University of Science and Technology) for valuable comments and discussions that improved this work.

## Author contributions

S.K.Z. and P.X. conceived the idea and supervised the study. H.Z. designed and implemented the framework, conducted the pre-training, structure-prediction and ncRNA-classification experiments, analysed the results and wrote the paper. R.L. conducted the zero-shot experiments, wrote the corresponding section and revised the overall writing. A.C. analysed the sequence-similarity results and the geometric and topological correctness of the predicted structures. H.C. processed sequence-similarity data for the training and test datasets and visualized the CASP15 and CASP16 datasets. F.Z. conducted the IRES-identification experiments. F.T. improved the overview of structRFM. T.Y. visualized the tertiary structures. X.L. designed the experiment evaluating the model across various sequence lengths. Y.G. improved the t-SNE visualization. All authors read the paper, contributed to the discussion and approved the final version.

## Additional information

**Supplementary information.** The Supplementary Information for this paper is publicly available.

**Correspondence and requests for materials.** should be addressed to Peng Xiong or S. Kevin Zhou.

---

**Extended Data Fig. 1:**
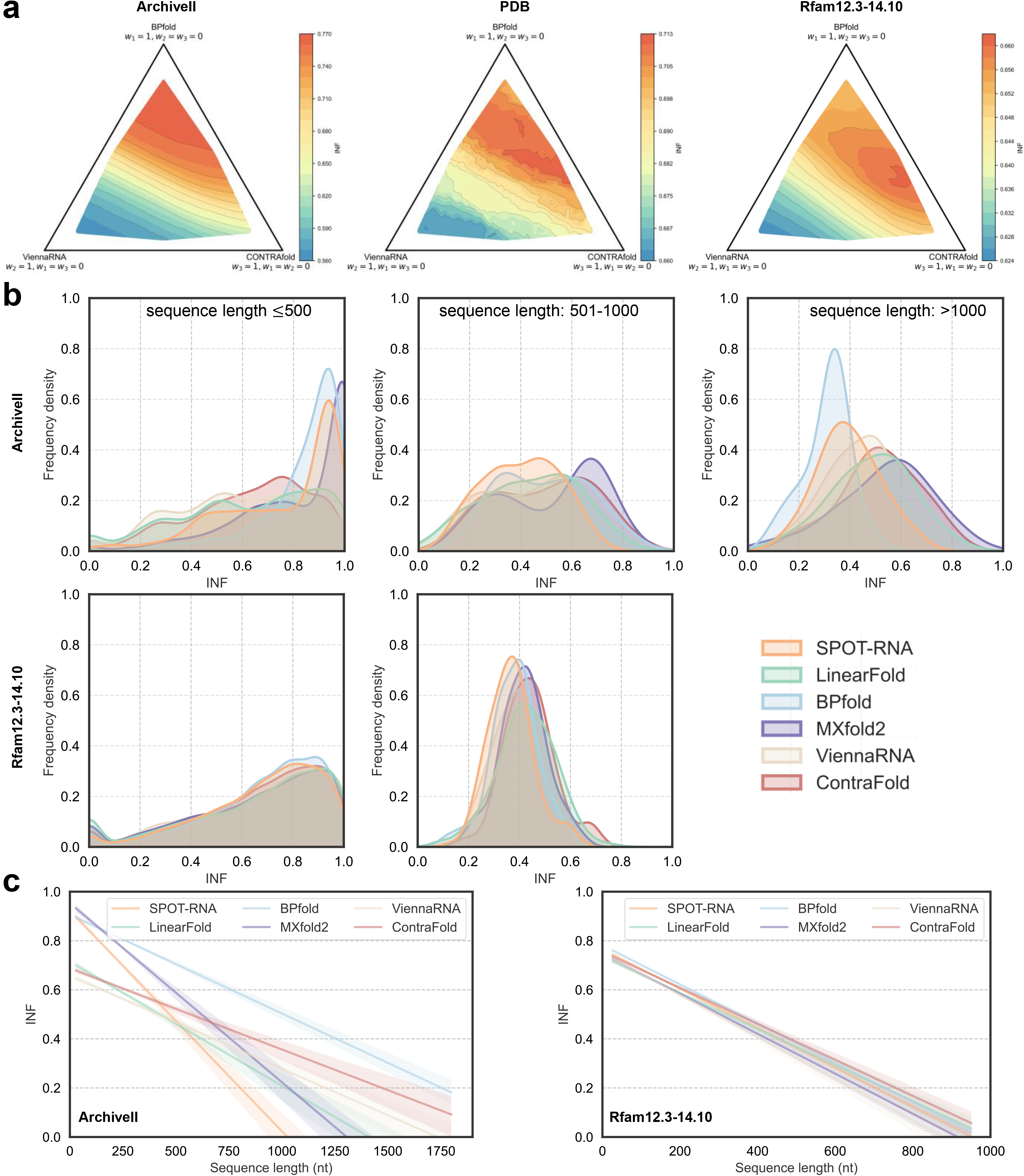
MUSES strategy for RNA secondary structure annotation and benchmarking of existing prediction methods across variable sequence lengths. **a** Ternary simplex visualization of the MUSES performance landscape across weighted combination configurations. Each vertex corresponds to a single standalone method (BPfold, ViennaRNA or CONTRAfold), and interior points represent weighted ensemble models. We normalized weighting coefficients for proportional equivalence and applied linear interpolation over a dense grid to smooth the INF metric distribution. Colour gradients (blue to red) represent INF values, with contour lines for enhanced resolution. This visualization illustrates how ensemble weighting modulates MUSES performance for robust secondary-structure annotation and identifies optimal weight-coefficient regimes with associated trade-offs. **b** Benchmark evaluation of six established multi-source RNA secondary-structure prediction methods [22–25, 37, 44, 45, 78] on the Rfam [66, 67] and full ArchiveII [26] (*n* = 3,975 independent RNA sequences) datasets, stratified by three sequence-length categories (< 500 nt, 501–1000 nt, > 1000 nt). **c** Linear fits extrapolating prediction performance for the six methods on both datasets. Lines are linear regression fits, and shaded bands represent 95% confidence intervals. A substantial performance degradation is observed with increasing sequence length, highlighting the current challenge of modelling RNA secondary structures for long RNA sequences.

**Extended Data Fig. 2:**
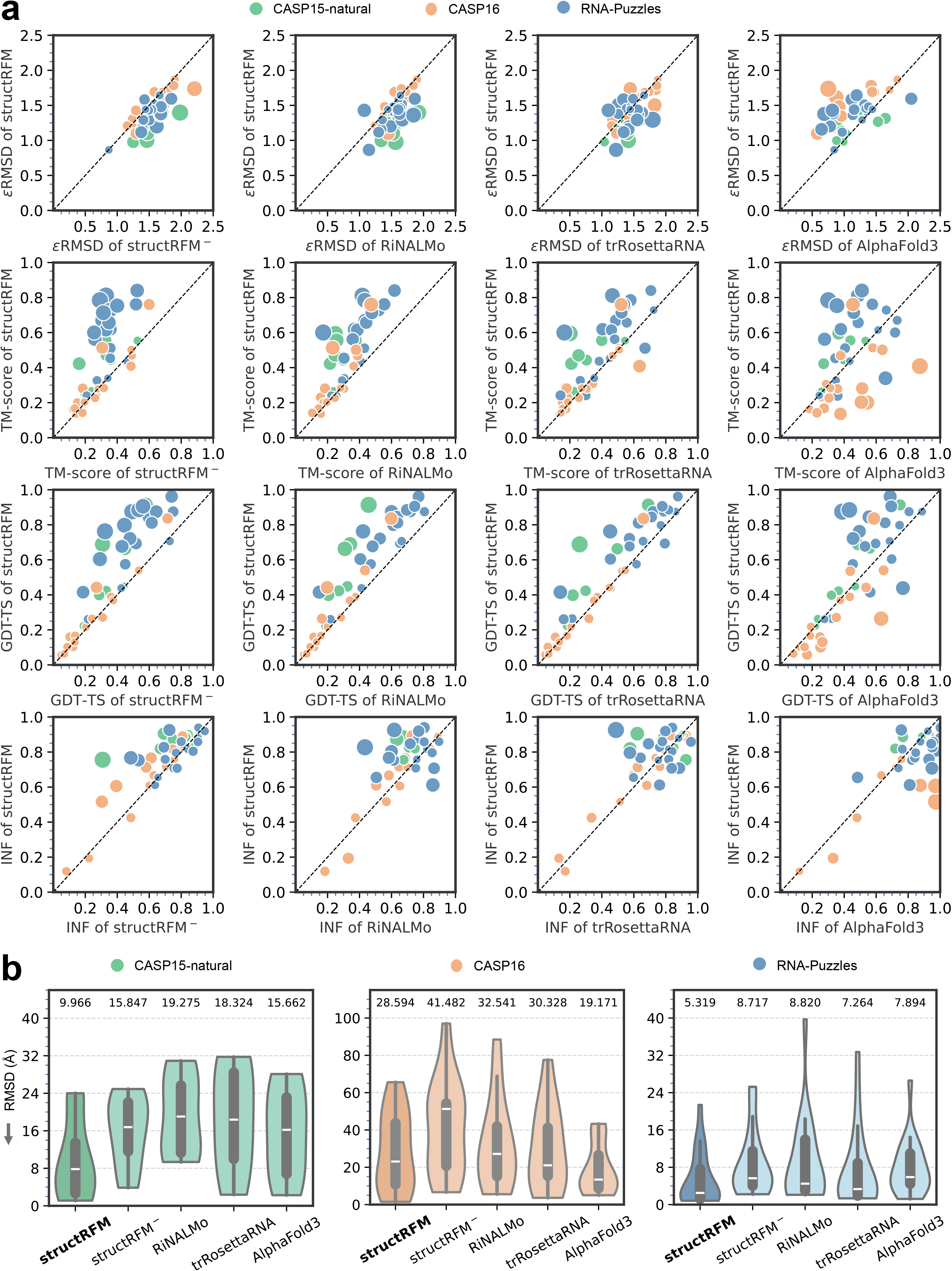
Detailed comparison of structRFM, structRFM*^−^*, RiNALMo-Mega [2], trRosettaRNA [28] and AlphaFold3 [31] for tertiary-structure prediction on the CASP15-natural [29] (8 RNAs), CASP16 (13 RNAs) [30] and RNA-Puzzles [48] (20 RNAs) datasets. **a** Head-to-head comparison of structRFM with other models using *ɛ*RMSD [72], TM-score [73], GDT-TS [74] and INF [75]. Circle size indicates the absolute difference between the two methods. **b** Violin plots of RMSD values. For each method in **b**, *n* = 8 independent RNA targets for CASP15-natural, *n* = 13 independent RNA targets for CASP16, and *n* = 20 independent RNA targets for RNA-Puzzles. The inner box plots show the median (white centre line), 25th and 75th percentiles (lower and upper bounds of the thick line), and minimum and maximum (lower and upper thin whiskers).

**Extended Data Fig. 3:**
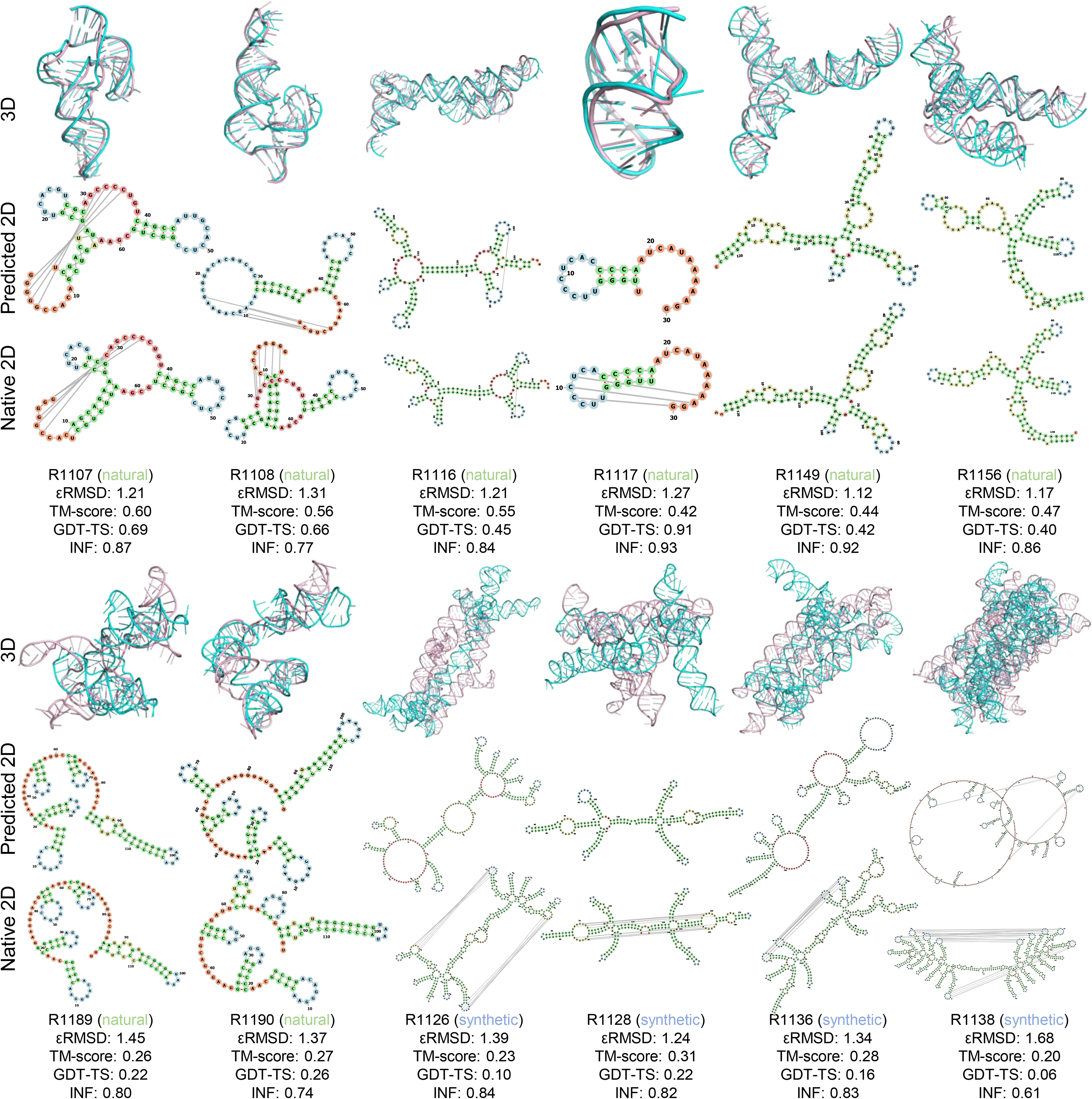
Visualization of structures predicted by structRFM (Zfold) and native structures in the CASP15 dataset. The *ɛ*RMSD, TM-score and GDT-TS of tertiary structures, together with the INF of secondary structures extracted from tertiary structures, are annotated. In the secondary-structure diagrams, stems are shown in green, multiple loops in red, interior loops in yellow, hairpin loops in blue, and 5’ and 3’ unpaired regions in orange. In general, structRFM performs better on natural targets than on synthetic targets containing dense long-range interactions, but sometimes fails to predict short local stems (in green), resulting in large multiple loops (in red).

**Extended Data Fig. 4:**
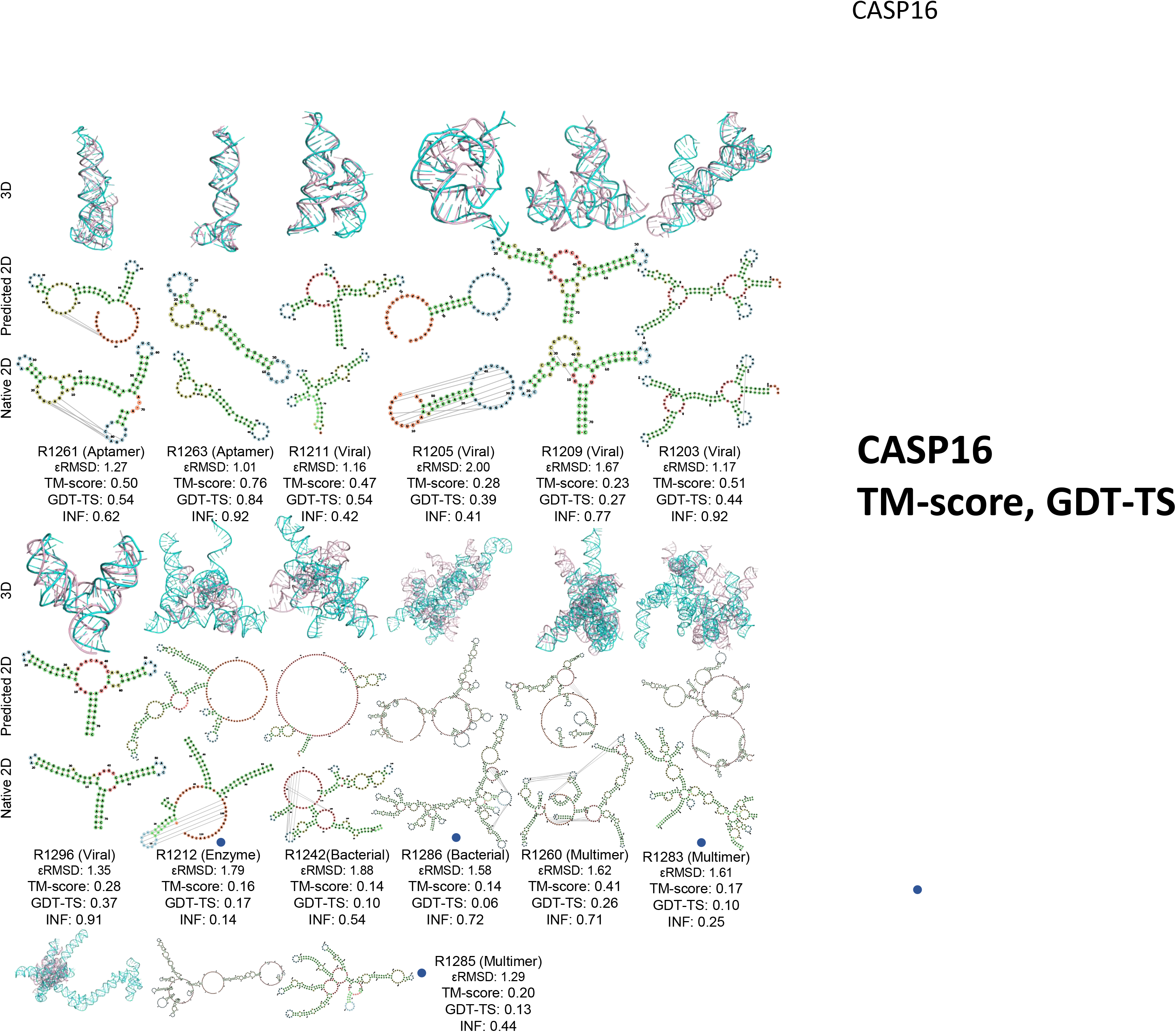
Visualization of structures predicted by structRFM (Zfold) and native structures in the CASP16 dataset. The *ɛ*RMSD, TM-score and GDT-TS of tertiary structures, together with the INF of secondary structures extracted from tertiary structures, are annotated. In the secondary-structure diagrams, stems are shown in green, multiple loops in red, interior loops in yellow, hairpin loops in blue, and 5’ and 3’ unpaired regions in orange. In general, structRFM performs better on viral and aptamer targets but performs worse on multimer and bacterial targets. For targets marked with blue circles, the sequences used for CASP16 prediction differ substantially from those of the PDB ground-truth structures because regions unresolved in the experimental structures were not modelled.

**Extended Data Fig. 5:**
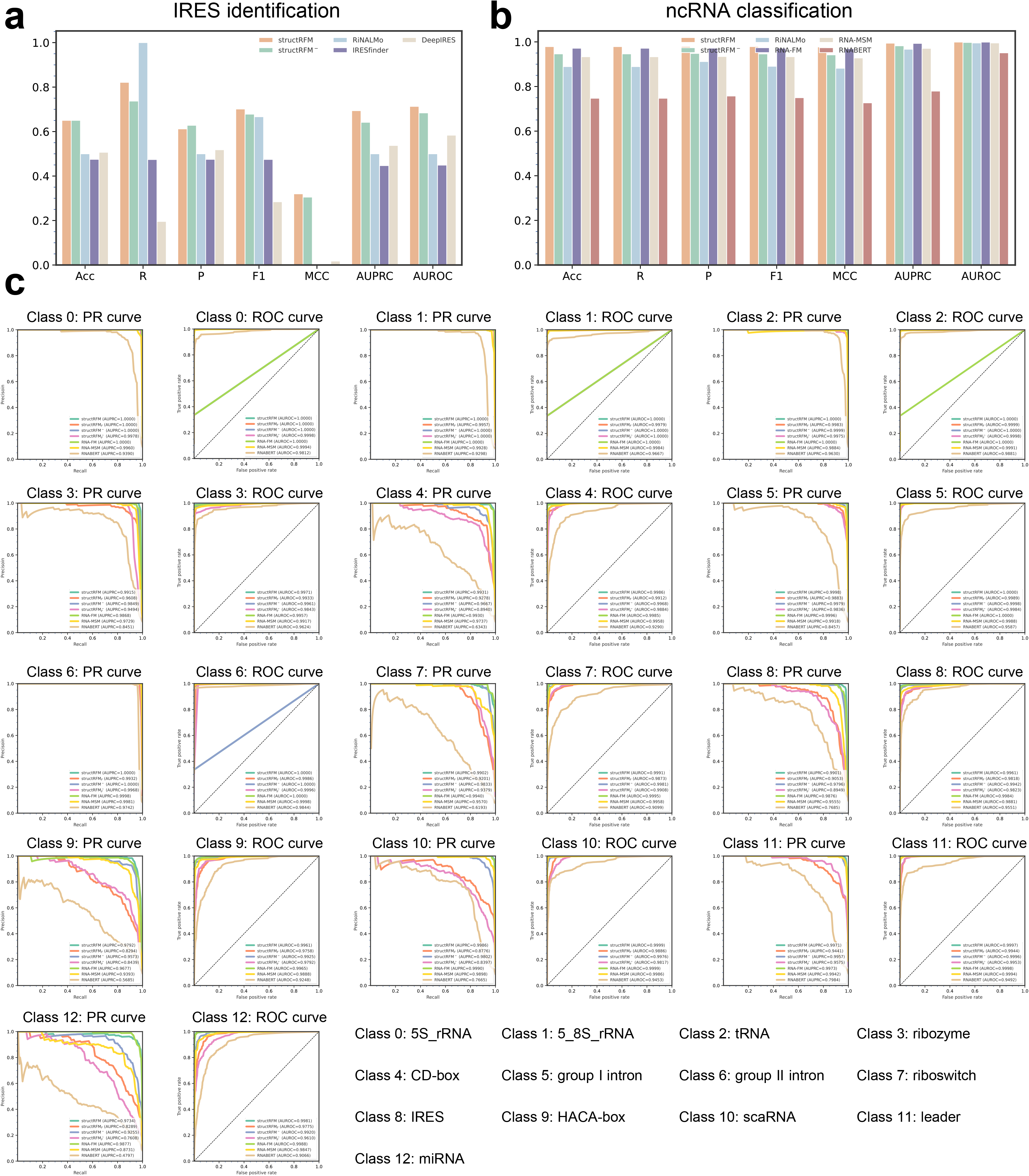
Detailed visualization of IRES identification and ncRNA classification. **a** Overall comparison of methods for IRES identification using seven metrics. **b** Overall comparison of methods for ncRNA classification using seven metrics. **c** PR and ROC curves for the 13 individual classes in the nRC dataset.

**Extended Data Fig. 6:**
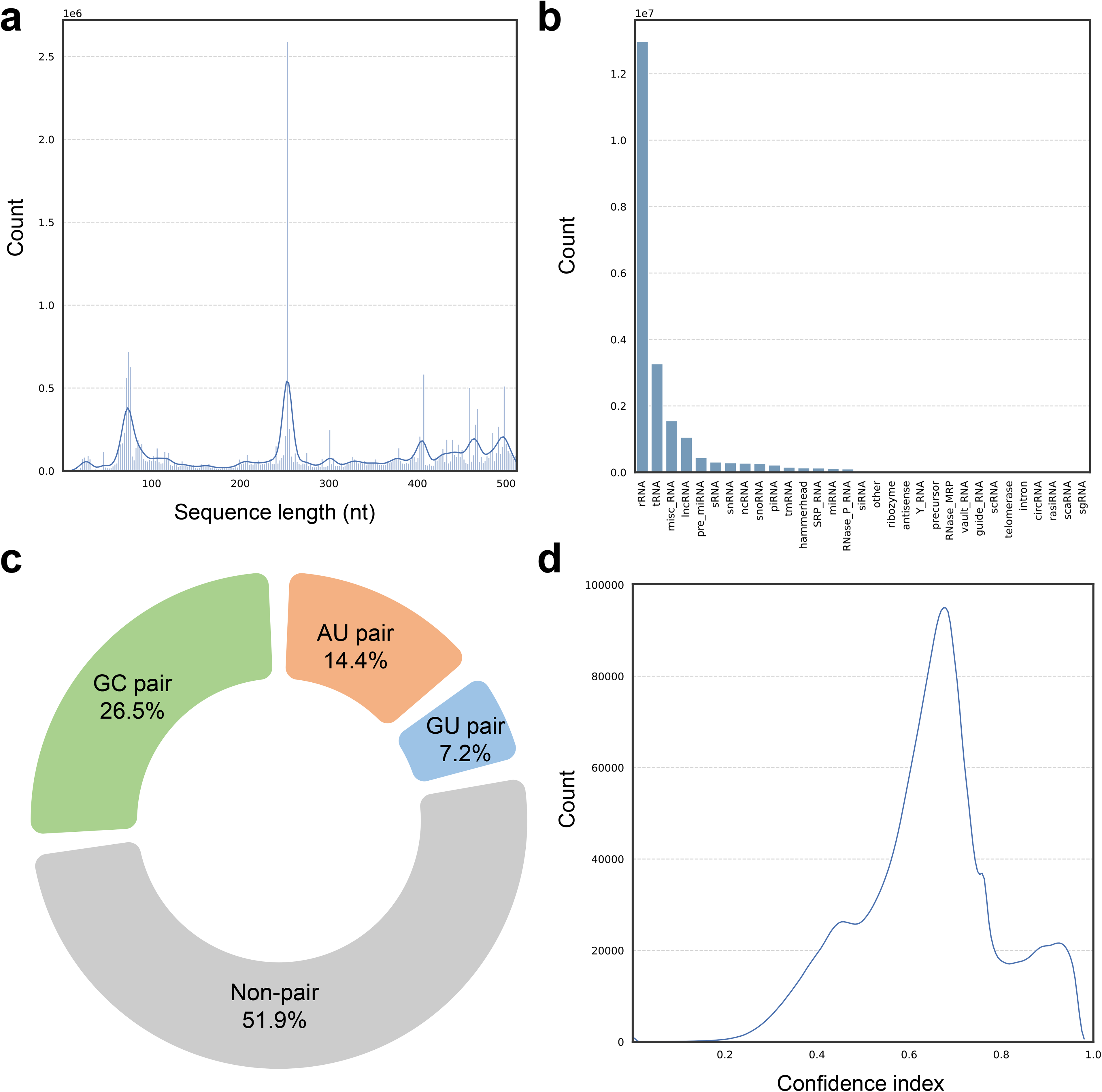
Overall statistics of our constructed pre-training sequence–structure dataset. **a** Distribution of sequence lengths, with most RNA sequences centred around 250 nucleotides. **b** Number and distribution of RNA families (categories). rRNA and tRNA dominate the dataset, accounting for 66.3% and 16.7%, respectively. **c** Composition of base-pairing interactions in secondary structures. Canonical base pairs represent approximately half of all interactions, including GC (26.5%), AU (14.4%) and GU (7.2%) pairs. **d** Distribution of confidence scores for predicted secondary structures. Most predictions have high confidence scores, supporting the reliability of the base-pairing annotations.

**Extended Data Table 1:**
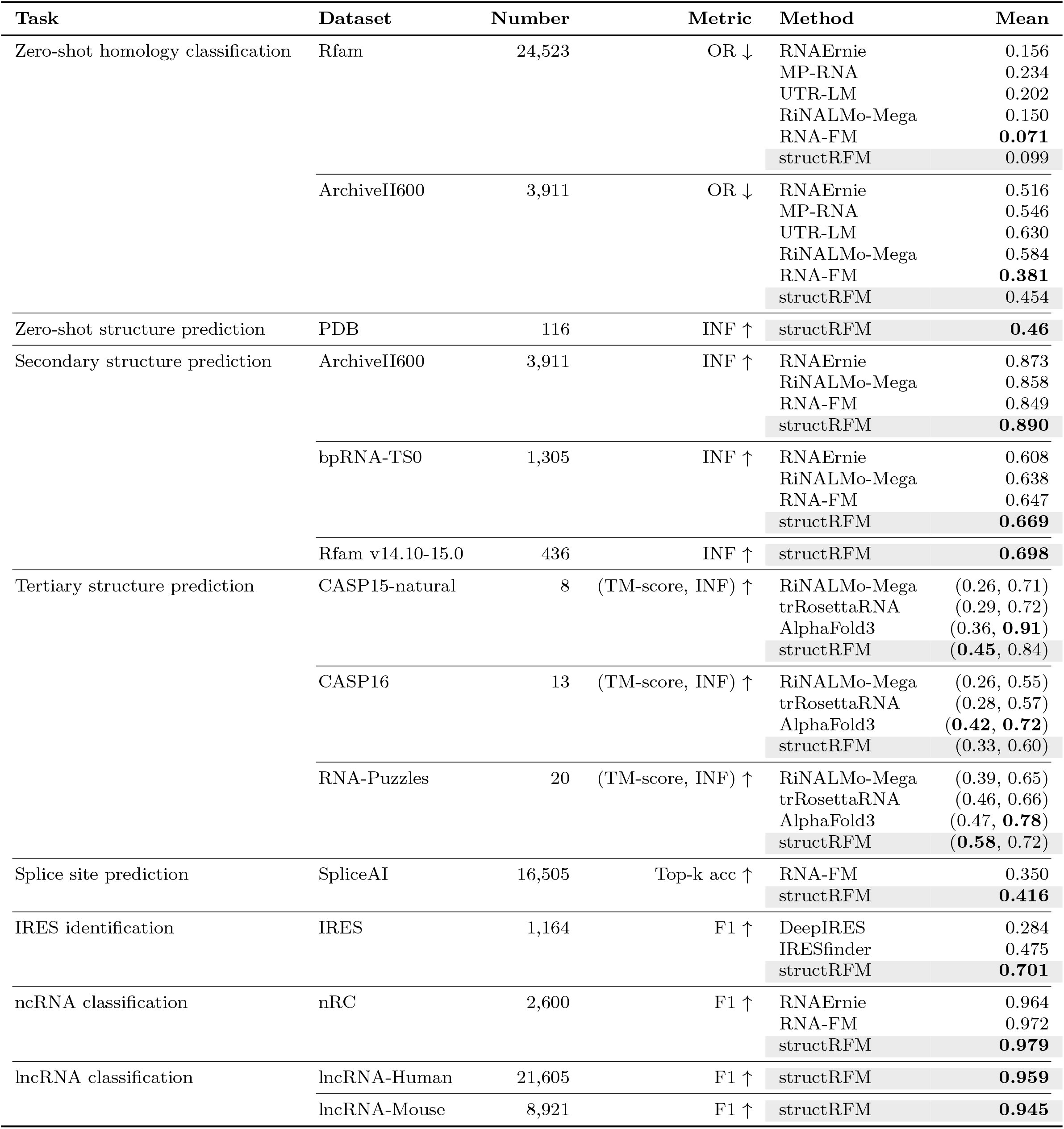
Overall performance of evaluating structRFM on zero-shot, structure, and function inference tasks.

**Extended Data Table 2:** Summary of used datasets.

| **Dataset** | **Task** | **Phase** | **# Sequences** | **Length** |
| --- | --- | --- | --- | --- |
| RNAcentral [41] | - | pre-train | 21,477,078 | $\leq 512$ |
| Rfam [66, 67] | Zero-shot homology classification | test | 24,523 | 19-4,002 |
| ArchiveII600 [37] | Zero-shot homology classification | test | 3,911 | 28-600 |
| PDB [46] | Zero-shot secondary structure | test | 116 | 32-355 |
| RNAStrAlign [35] | Secondary structure prediction | train | 19,313 | 30-600 |
| bpRNA-TR0 [36] | Secondary structure prediction | train | 12,114 | 33-498 |
| bpRNA-TS0 [36] | Secondary structure prediction | test | 1,305 | 22-499 |
| ArchiveII600 [37] | Secondary structure prediction | test | 3,911 | 28-600 |
| ArchiveII [37] | Secondary structure method benchmarking | test | 3,975 | 28-2,968 |
| Rfam14.10-15.0 [66] | Secondary structure prediction | test | 436 | 26-210 |
| PDB [46] | Tertiary structure prediction | train | 3,633 | 11-960 |
| CASP15 [27] | Tertiary structure prediction | test | 12 | 30-720 |
| CASP16 [28] | Tertiary structure prediction | test | 13 | 59-580 |
| RNA-Puzzles [48, 68] | Tertiary structure prediction | test | 20 | 37-189 |
| Splice [4] | Splice site prediction | train | 162,706 | 100 |
| Splice [4] | Splice site prediction | test | 16,505 | 100 |
| IRES [29, 30] | IRES identification | train | 12,998 | 174 |
| IRES [29, 30] | IRES identification | test | 1,164 | 174 |
| nRC [31] | ncRNA classification | train | 6,320 | 38-1,136 |
| nRC [31] | ncRNA classification | test | 2,600 | 39-1,182 |
| lncRNA-Human [56] | lncRNA classification | train | 72,768 | 9-3000 |
| lncRNA-Human [56] | lncRNA classification | test | 21,605 | 10-3000 |
| lncRNA-Mouse [56] | lncRNA classification | train | 29,625 | 8-3000 |
| lncRNA-Mouse [56] | lncRNA classification | test | 8,921 | 43-3000 |

**Extended Data Table 3:** Comparative overview of structRFM with other RNA language models. Seq. is short for sequence and Stru. is short for secondary structure. The performances of overlap ratio (OR, lower is better) for zero-shot homology classification on Rfam [66, 67] dataset are displayed.

| Model name | Model size(M) | Pre-training dataset | Structure information | Task-agnostic pre-training | Zero-shot (OR↓) |
| --- | --- | --- | --- | --- | --- |
| RiNALMo-Micro | 33 | ~36M Seq. | No | Yes | 0.167 |
| RiNALMo-Mega | 148 | ~36M Seq. | No | Yes | 0.150 |
| RiNALMo-Giga | 651 | ~36M Seq. | No | Yes | 0.115 |
| RNAErnie | 86 | ~23M Seq. | No | Yes | 0.156 |
| BiRNA-BERT | 116 | ~37M Seq | No | Yes | 0.207 |
| AIDO.RNA | 650 | ~42M Seq. | No | Yes | 0.102 |
| AIDO.RNA | 1,600 | ~42M Seq. | No | Yes | 0.110 |
| SpliceBERT | 20 | ~2M Seq. | No | Yes | 0.239 |
| RNA-FM | 100 | ~23M Seq. | No | Yes | 0.071 |
| ERNIE-RNA | 86 | ~20M Seq. | Pairwise score | Yes | 0.090 |
| MP-RNA | 186 | - | ViennaRNA | No | 0.234 |
| UTR_LM | 1 | ~2.3M Seq., Stru. | ViennaRNA | No | 0.202 |
| structRFM | 86 | ~21M Seq., Stru. | MUSES | Yes | 0.099 |

## References

[1] Chen, J., et al. Interpretable rna foundation model from unannotated data for highly accurate rna structure and function predictions. arXiv preprint arXiv:2204.00300 (2022).

[2] Penić, R. J., Vlašić, T., Huber, R. G., Wan, Y. & Šikić, M. Rinalmo: General-purpose rna language models can generalize well on structure prediction tasks. Nature Communications 16, 5671 (2025).

[3] Cao, X., Zhang, Y., Ding, Y. & Wan, Y. Identification of rna structures and their roles in rna functions. Nature Reviews Molecular Cell Biology 25, 784–801 (2024).

[4] Mortimer, S. A., Kidwell, M. A. & Doudna, J. A. Insights into rna structure and function from genome-wide studies. Nature Reviews Genetics 15, 469–479 (2014).

[5] Cooper, T. A., Wan, L. & Dreyfuss, G. Rna and disease. Cell 136, 777–793 (2009).

[6] Fan, X., Yang, Y., Chen, C. & Wang, Z. Pervasive translation of circular rnas driven by short ires-like elements. Nature Communications 13, 3751 (2022).

[7] Jaganathan, K. et al. Predicting splicing from primary sequence with deep learning. Cell 176, 535–548 (2019).

[8] Ozsolak, F. & Milos, P. M. Rna sequencing: advances, challenges and opportunities. Nature reviews genetics 12, 87–98 (2011).

[9] He, Y. et al. Generalized biological foundation model with unified nucleic acid and protein language. Nature Machine Intelligence 7, 942–953 (2025). URL 10.1038/s42256-025-01044-4.

[10] Brixi, G. et al. Genome modelling and design across all domains of life with Evo 2. Nature 652, 1349–1361 (2026).

[11] Li, Y., Feng, C., Zhang, X. & Zhang, Y. Ab initio rna structure prediction with composite language model and denoised end-to-end learning. bioRxiv 2025–03 (2025).

[12] Shen, T. et al. Accurate rna 3d structure prediction using a language model-based deep learning approach. Nature Methods 1–12 (2024).

[13] Chu, Y. et al. A 5’ utr language model for decoding untranslated regions of mrna and function predictions. Nature Machine Intelligence 6, 449–460 (2024).

[14] Chen, K. et al. Self-supervised learning on millions of primary rna sequences from 72 vertebrates improves sequence-based rna splicing prediction. Briefings in bioinformatics 25, bbae163 (2024).

[15] Wang, H., Zhang, Y., Chen, J., Zhan, J. & Zhou, Y. Zero-shot benchmarking of rna language models in structural, functional, and evolutionary learning. Briefings in Bioinformatics 27, bbag098 (2026). URL 10.1093/bib/bbag098.

[16] Wang, N. et al. Multi-purpose rna language modelling with motif-aware pretraining and type-guided fine-tuning. Nature Machine Intelligence 6, 548–557 (2024).

[17] Zou, S., et al. A large-scale foundation model for RNA function and structure prediction (2024).

[18] Yu, H. et al. An interpretable rna foundation model for exploring functional rna motifs in plants. Nature Machine Intelligence 6, 1616–1625 (2024).

[19] Akiyama, M. & Sakakibara, Y. Informative rna base embedding for rna structural alignment and clustering by deep representation learning. NAR genomics and bioinformatics 4, lqac012 (2022).

[20] Yin, W. et al. Ernie-rna: an rna language model with structure-enhanced representations. Nature Communications 16, 10076 (2025).

[21] Yang, H. & Li, K. Al-Onaizan, Y., Bansal, M. & Chen, Y. (eds) *MP-RNA: unleashing multi-species RNA foundation model via calibrated secondary structure prediction*. (eds Al-Onaizan, Y., Bansal, M. & Chen, Y.) Findings of the Association for Computational Linguistics: EMNLP 2024, Miami, Florida, USA, November 12-16, 2024, 5278–5296 (Association for Computational Linguistics, 2024).

[22] Hofacker, I. L. et al. Fast folding and comparison of rna secondary structures. Monatshefte fur chemie 125, 167–167 (1994).

[23] Lorenz, R. et al. Viennarna package 2.0. Algorithms for molecular biology 6, 1–14 (2011).

[24] Do, C. B., Woods, D. A. & Batzoglou, S. Contrafold: Rna secondary structure prediction without physics-based models. Bioinformatics 22, e90–e98 (2006).

[25] Zhu, H. et al. Deep generalizable prediction of RNA secondary structure via base pair motif energy. Nature Communications 16, 5856 (2025). URL 10.1038/s41467-025-60048-1.

[26] Sloma, M. F. & Mathews, D. H. Exact calculation of loop formation probability identifies folding motifs in rna secondary structures. RNA 22, 1808–1818 (2016).

[27] Danaee, P. et al. bprna: large-scale automated annotation and analysis of rna secondary structure. Nucleic acids research 46, 5381–5394 (2018).

[28] Wang, W. et al. trrosettarna: automated prediction of rna 3d structure with transformer network. Nature communications 14, 7266 (2023).

[29] Das, R. et al. Assessment of three-dimensional rna structure prediction in casp15. *Proteins: Structure*, Function, and Bioinformatics 91, 1747–1770 (2023).

[30] Kretsch, R. C. et al. Assessment of nucleic acid structure prediction in casp16. *Proteins: Structure*, Function, and Bioinformatics (2025).

[31] Abramson, J. et al. Accurate structure prediction of biomolecular interactions with alphafold 3. Nature 630, 493–500 (2024).

[32] Chen, C.-K. et al. Structured elements drive extensive circular rna translation. Molecular Cell 81, 4300–4318.e13 (2021).

[33] Weingarten-Gabbay, S. et al. Systematic discovery of cap-independent translation sequences in human and viral genomes. Science 351, aad4939 (2016).

[34] Fiannaca, A., La Rosa, M., La Paglia, L., Rizzo, R. & Urso, A. nrc: non-coding rna classifier based on structural features. BioData mining 10, 1–18 (2017).

[35] Van der Maaten, L. & Hinton, G. Visualizing data using t-sne. Journal of machine learning research 9, 2579–2605 (2008).

[36] Zhang, Y. et al. Multiple sequence alignment-based rna language model and its application to structural inference. Nucleic Acids Research 52, e3–e3 (2024).

[37] Sato, K., Akiyama, M. & Sakakibara, Y. Rna secondary structure prediction using deep learning with thermodynamic integration. Nature communications 12, 941 (2021).

[38] Tan, Z., Fu, Y., Sharma, G. & Mathews, D. H. Turbofold ii: Rna structural alignment and secondary structure prediction informed by multiple homologs. Nucleic acids research 45, 11570–11581 (2017).

[39] Zakov, S., Goldberg, Y., Elhadad, M. & Ziv-Ukelson, M. Rich parameterization improves rna structure prediction. Journal of Computational Biology 18, 1525– 1542 (2011).

[40] Zhang, C., Zhang, Y. & Pyle, A. M. rmsa: a sequence search and alignment algorithm to improve rna structure modeling. Journal of Molecular Biology 435, 167904 (2023).

[41] Altschul, S. F., Gish, W., Miller, W., Myers, E. W. & Lipman, D. J. Basic local alignment search tool. Journal of molecular biology 215, 403–410 (1990).

[42] Sweeney, B. A. et al. Rnacentral 2021: secondary structure integration, improved sequence search and new member databases. Nucleic Acids Res. 49, D212–D220 (2021).

[43] Coordinators, N. R. Database resources of the national center for biotechnology information. Nucleic acids research 41, D8–D20 (2012).

[44] Singh, J., Hanson, J., Paliwal, K. & Zhou, Y. Rna secondary structure prediction using an ensemble of two-dimensional deep neural networks and transfer learning. Nature communications 10, 5407 (2019).

[45] Singh, J. et al. Improved rna secondary structure and tertiary base-pairing prediction using evolutionary profile, mutational coupling and two-dimensional transfer learning. Bioinformatics 37, 2589–2600 (2021).

[46] Berman, H. M. et al. The protein data bank. Nucleic acids research 28, 235–242 (2000).

[47] Rose, P. W. et al. The rcsb protein data bank: integrative view of protein, gene and 3d structural information. Nucleic acids research gkw1000 (2016).

[48] Cruz, J. A. et al. Rna-puzzles: a casp-like evaluation of rna three-dimensional structure prediction. Rna 18, 610–625 (2012).

[49] Popenda, M. et al. Entanglements of structure elements revealed in rna 3d models. Nucleic acids research 49, 9625–9632 (2021).

[50] Gren, B. A., Antczak, M., Zok, T., Sulkowska, J. I. & Szachniuk, M. Knotted artifacts in predicted 3d rna structures. PLOS Computational Biology 20, e1011959 (2024).

[51] Carrascoza, F., Antczak, M., Miao, Z., Westhof, E. & Szachniuk, M. Evaluation of the stereochemical quality of predicted rna 3d models in the rna-puzzles submissions. Rna 28, 250–262 (2022).

[52] Ren, Y. et al. Beacon: Benchmark for comprehensive rna tasks and language models. Advances in Neural Information Processing Systems 37, 92891–92921 (2024).

[53] Zhao, J. et al. Iresfinder: Identifying rna internal ribosome entry site in eukaryotic cell using framed k-mer features. Journal of genetics and genomics= Yi chuan xue bao 45, 403–406 (2018).

[54] Zhao, J. et al. Deepires: a hybrid deep learning model for accurate identification of internal ribosome entry sites in cellular and viral mrnas. Briefings in Bioinformatics 25, bbae439 (2024).

[55] Chantsalnyam, T., Siraj, A., Tayara, H. & Chong, K. T. ncrdense: a novel computational approach for classification of non-coding rna family by deep learning. Genomics 113, 3030–3038 (2021).

[56] Frankish, A. et al. GENCODE 2021. Nucleic acids research 49, D916–D923 (2021).

[57] Vaswani, A. et al. Guyon, I. et al. (eds) *Attention is all you need*. (eds Guyon, I. et al.) In 31th Conference on Neural Information Processing Systems, 5998–6008 (NeurIPS, 2017).

[58] Gu, A. & Dao, T. Mamba: Linear-time sequence modeling with selective state spaces. arXiv preprint arXiv:2312.00752 (2023).

## Methods references

[59] Zhang, Z. et al. Rnagenesis: Foundation model for enhanced rna sequence generation and structural insights. bioRxiv 2024–12 (2024).

[60] Fu, L. et al. Ufold: fast and accurate rna secondary structure prediction with deep learning. Nucleic acids research 50, e14–e14 (2022).

[61] Chen, X., Li, Y., Umarov, R., Gao, X. & Song, L. Wallach, H. M. et al. (eds) *RNA secondary structure prediction by learning unrolled algorithms*. (eds Wallach, H. M. et al.) 8th International Conference on Learning Representations, ICLR 2020, Addis Ababa, Ethiopia, April 26-30, 2020, 8024–8035 (OpenReview.net, 2020). URL https://openreview.net/forum?id=S1eALyrYDH.

[62] Zuker, M. & Stiegler, P. Optimal computer folding of large rna sequences using thermodynamics and auxiliary information. Nucleic acids research 9, 133–148 (1981).

[63] Chaudhury, S., Lyskov, S. & Gray, J. J. Pyrosetta: a script-based interface for implementing molecular modeling algorithms using rosetta. Bioinformatics 26, 689–691 (2010).

[64] Devlin, J., Chang, M., Lee, K. & Toutanova, K. Burstein, J., Doran, C. & Solorio, T. (eds) *BERT: pre-training of deep bidirectional transformers for language understanding*. (eds Burstein, J., Doran, C. & Solorio, T.) Proceedings of the 2019 Conference of the North American Chapter of the Association for Computational Linguistics: Human Language Technologies, 4171–4186 (Association for Computational Linguistics, 2019).

[65] Paszke, A. et al. Wallach, H. M. et al. (eds) Pytorch: An imperative style, high-performance deep learning library. (eds Wallach, H. M. et al.) Advances in Neural Information Processing Systems 32: Annual Conference on Neural Information Processing Systems 2019, NeurIPS 2019, December 8-14, 2019, Vancouver, BC, Canada, 8024–8035 (2019). URL https://proceedings.neurips.cc/paper/2019/hash/bdbca288fee7f92f2bfa9f7012727740-Abstract.html.

[66] Kalvari, I. et al. Non-coding rna analysis using the rfam database. Current protocols in bioinformatics 62, e51 (2018).

[67] Kalvari, I. et al. Rfam 14: expanded coverage of metagenomic, viral and microrna families. Nucleic acids research 49, D192–D200 (2021).

[68] Miao, Z. et al. Rna-puzzles round iv: 3d structure predictions of four ribozymes and two aptamers. Rna 26, 982–995 (2020).

[69] Yang, H. et al. Tools for the automatic identification and classification of rna base pairs. Nucleic Acids Research 31, 3450–3460 (2003).

[70] Sweeney, B. A. et al. R2dt is a framework for predicting and visualising rna secondary structure using templates. Nature Communications 12, 3494 (2021).

[71] Kerpedjiev, P., Hammer, S. & Hofacker, I. L. Forna (force-directed rna): Simple and effective online rna secondary structure diagrams. Bioinformatics 31, 3377– 3379 (2015).

[72] Bottaro, S., Di Palma, F. & Bussi, G. The role of nucleobase interactions in rna structure and dynamics. Nucleic acids research 42, 13306–13314 (2014).

[73] Xu, J. & Zhang, Y. How significant is a protein structure similarity with tm-score= 0.5? Bioinformatics 26, 889–895 (2010).

[74] Zemla, A. Lga: a method for finding 3d similarities in protein structures. Nucleic acids research 31, 3370–3374 (2003).

[75] Parisien, M., Cruz, J. A., Westhof, E. & Major, F. New metrics for comparing and assessing discrepancies between rna 3d structures and models. Rna 15, 1875–1885 (2009).

[76] Zhu, H. structRFM pre-training data (2026). URL 10.5281/zenodo.18272040.

[77] Zhu, H. heqin-zhu/structrfm: v0.0.8 (2026). URL 10.5281/zenodo.18274292.

[78] Huang, L. et al. Linearfold: linear-time approximate rna folding by 5’-to-3’dynamic programming and beam search. Bioinformatics 35, i295–i304 (2019).

