## Supplementary for "An open structure-guided RNA foundation model for generalizable prediction"

### 1 Supplementary Methods

The main Methods describe the pre-training corpus, model architecture, SgMLM objective and downstream task formulations. To avoid duplicating those descriptions, this section reports only additional fine-tuning settings and the controlled comparison between BPfold- and MUSES-derived pre-training annotations.

#### 1.1 Additional fine-tuning hyperparameters

All downstream experiments were initialized from the same pre-trained checkpoint unless otherwise stated. IRES-identification models were fine-tuned for 100 epochs with a batch size of 128 and a learning rate of $2 \times 10^{-3}$. Zfold used a batch size of 1 and an initial learning rate of $5 \times 10^{-4}$. For ncRNA classification, the batch size was 16 and the learning rate was $1 \times 10^{-4}$. Secondary-structure prediction used a batch size of 1 and a learning rate of $5 \times 10^{-4}$, whereas splice-site prediction used a batch size of 32 and a learning rate of $3 \times 10^{-5}$.

#### 1.2 Comparison of MUSES- and BPfold-derived annotations

To isolate the effect of the pre-training structure annotations, we compared structRFM models trained with MUSES- or BPfold-derived structures (Supplementary Table 9). Here ArchiveII600 denotes the 3,911-sequence subset of ArchiveII obtained by filtering sequences to at most 600 nt. For zero-shot homology classification, MUSES gives an OR of 0.454 on ArchiveII600, compared with 0.483 for BPfold, whereas the corresponding Rfam values are 0.099 and 0.097. For zero-shot structure prediction on the PDB set, MUSES increases INF from 0.44 to 0.46. The differences on the three function-prediction tasks are small and mixed.

The advantage of MUSES is more consistent on structure-prediction tasks. Relative to BPfold-derived annotations, MUSES increases INF from 0.873 to 0.890 on ArchiveII600 and from 0.641 to 0.669 on bpRNA-TS0. It also reduces RMSD from 14.8Å to 9.97Å on CASP15-natural and from 6.82Å to 5.32Å on RNA-Puzzles. Thus, the annotation ensemble provides its clearest benefit for structure-centric downstream inference rather than uniformly improving every task.

### 2 Supplementary figures

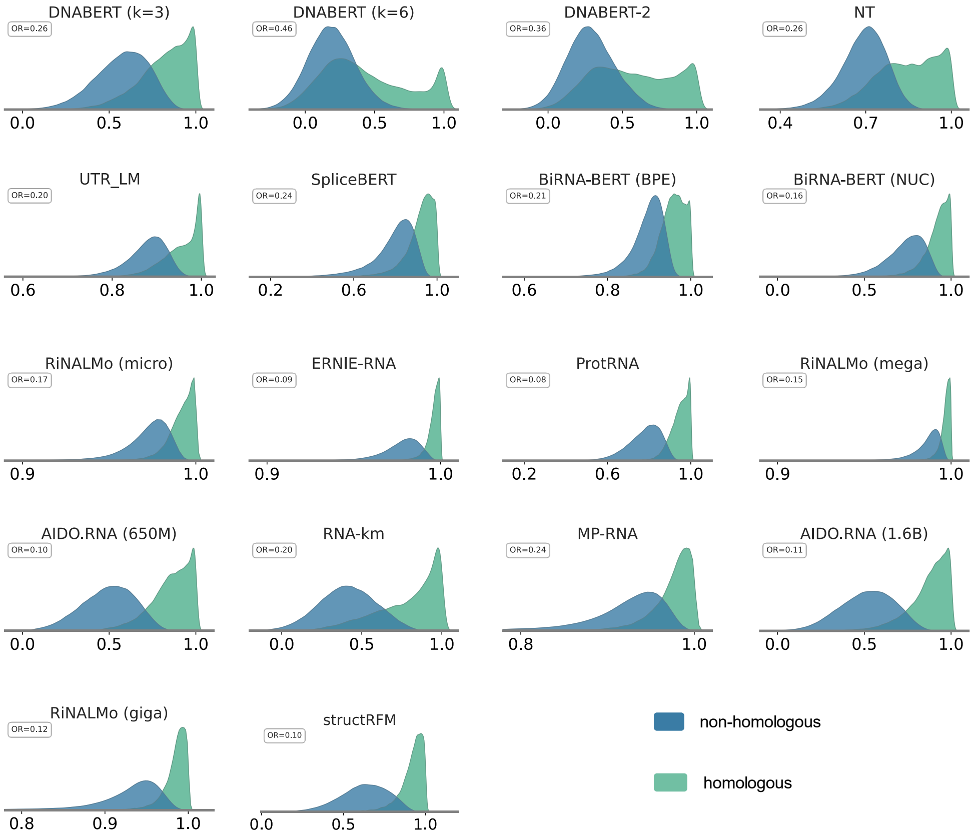


**Supplementary Fig. 1**: **Cosine-similarity distributions for homologous and non-homologous sequence pairs in Rfam.** The dataset contains 24,523 RNA sequences. Homologous pairs contain sequences from the same RNA family, whereas non-homologous pairs contain sequences from different families; each distribution comprises $n = 100{,}000$ sampled sequence pairs.

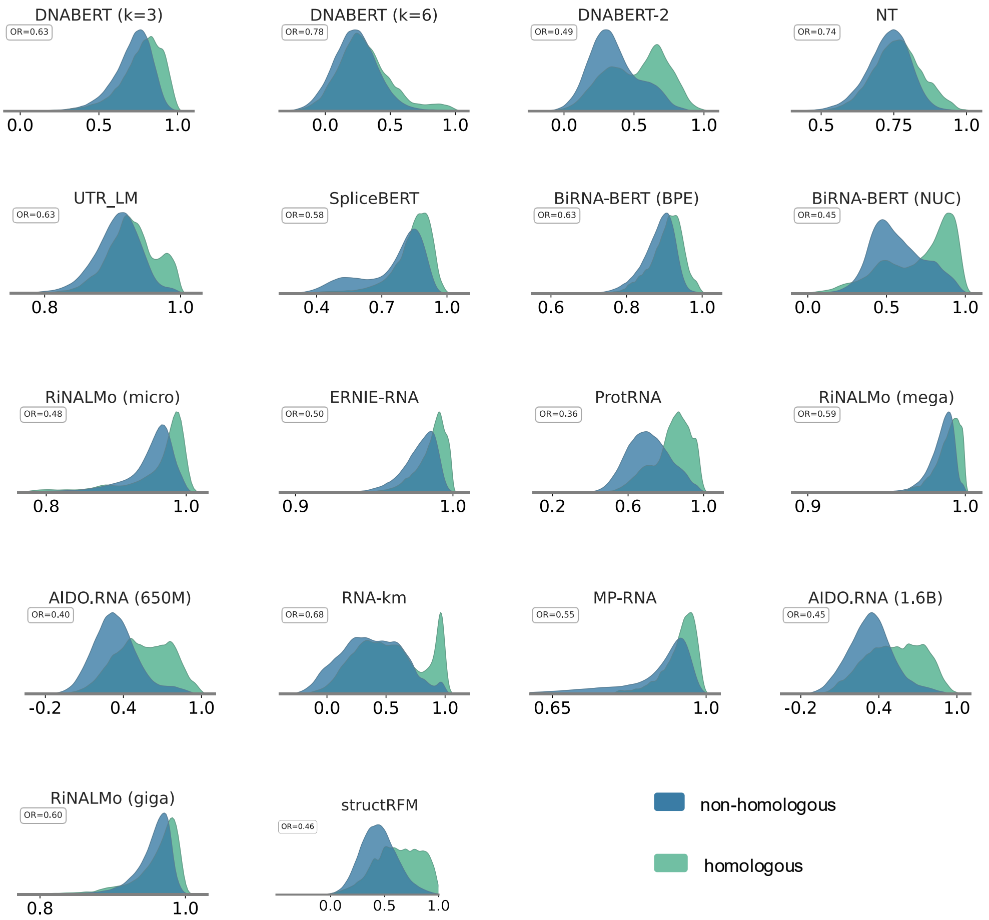


**Supplementary Fig. 2**: **Cosine-similarity distributions for homologous and non-homologous sequence pairs in ArchiveII600.** The dataset contains 3,911 RNA sequences. Homologous pairs contain sequences of the same RNA type, whereas non-homologous pairs contain sequences of different types; each distribution comprises $n = 100{,}000$ sampled sequence pairs.

#### 3 Supplementary tables

**Supplementary Table 1**: Secondary-structure prediction performance on ArchiveII600 (3,911 RNAs) and bpRNA-TS0 (1,305 RNAs). Results for the baseline methods are taken from RNAErnie [1].

| Method | ArchiveII600 | | | bpRNA-TS0 | | |
|---|---|---|---|---|---|---|
| | Precision | Recall | INF | Precision | Recall | INF |
| structRFM | 0.904 | 0.880 | 0.890 | 0.635 | 0.720 | 0.669 |
| structRFM$^{-}$ | 0.850 | 0.857 | 0.851 | 0.587 | 0.691 | 0.630 |
| RNABERT | 0.694 | 0.669 | 0.679 | 0.554 | 0.581 | 0.557 |
| RNA-MSM | 0.608 | 0.585 | 0.594 | 0.548 | 0.612 | 0.569 |
| RNA-FM | 0.857 | 0.844 | 0.849 | 0.596 | 0.716 | 0.647 |
| RNAstructure | 0.563 | 0.615 | 0.588 | 0.494 | 0.622 | 0.554 |
| RNAsoft | 0.665 | 0.594 | 0.628 | 0.497 | 0.626 | 0.558 |
| RNAfold | 0.565 | 0.627 | 0.595 | 0.494 | 0.631 | 0.558 |
| MXfold2 | 0.788 | 0.760 | 0.774 | 0.519 | 0.646 | 0.579 |
| Mfold | 0.428 | 0.383 | 0.405 | 0.501 | 0.627 | 0.560 |
| LinearFold | 0.641 | 0.617 | 0.629 | 0.561 | 0.581 | 0.51 |
| Eternafold | 0.667 | 0.622 | 0.644 | 0.516 | 0.666 | 0.586 |
| E2Efold | 0.738 | 0.665 | 0.701 | 0.140 | 0.129 | 0.134 |
| CONTRAFold | 0.607 | 0.679 | 0.642 | 0.528 | 0.655 | 0.588 |
| Contextfold | 0.873 | 0.821 | 0.847 | 0.529 | 0.607 | 0.567 |
| RNAErnie | 0.884 | 0.869 | 0.876 | 0.576 | 0.668 | 0.620 |

**Supplementary Table 2**: Family-level secondary-structure prediction performance of structRFM and structRFM$^{-}$ across the nine RNA families in ArchiveII600 (3,911 RNAs).

| Family | structRFM | | | structRFM$^{-}$ | | |
|---|---|---|---|---|---|---|
| | Precision | Recall | INF | Precision | Recall | INF |
| 16s | 0.684 | 0.688 | 0.683 | 0.725 | 0.718 | 0.717 |
| 23s | 0.332 | 0.295 | 0.311 | 0.552 | 0.487 | 0.516 |
| 5s | 0.975 | 0.985 | 0.980 | 0.962 | 0.971 | 0.966 |
| RNaseP | 0.848 | 0.825 | 0.834 | 0.818 | 0.793 | 0.802 |
| grp1 | 0.670 | 0.693 | 0.677 | 0.642 | 0.689 | 0.659 |
| srp | 0.794 | 0.821 | 0.803 | 0.752 | 0.800 | 0.771 |
| tRNA | 0.960 | 0.993 | 0.975 | 0.962 | 0.980 | 0.969 |
| telomerase | 0.578 | 0.619 | 0.595 | 0.435 | 0.475 | 0.453 |
| tmRNA | 0.810 | 0.703 | 0.748 | 0.737 | 0.670 | 0.698 |

**Supplementary Table 3**: Tertiary-structure prediction performance of fine-tuned structRFM (Zfold), structRFM$^{-}$, RiNALMo-Mega, trRosettaRNA and AlphaFold3 on CASP15 (12 RNAs: 8 natural and 4 synthetic), CASP16 (13 RNAs) and RNA-Puzzles (20 RNAs), evaluated using RMSD, $\epsilon$RMSD, TM-score, GDT-TS and INF.

| Model | Dataset | TM-score | GDT-TS | $\epsilon$RMSD | INF | RMSD |
|---|---|---|---|---|---|---|
| structRFM | RNA-Puzzles | 0.583 | 0.698 | 1.255 | 0.725 | 5.319 |
| | CASP15 | 0.382 | 0.380 | 1.313 | 0.820 | 20.130 |
| | CASP16 | 0.327 | 0.324 | 1.493 | 0.597 | 28.594 |
| structRFM$^{-}$ | RNA-Puzzles | 0.343 | 0.483 | 1.484 | 0.686 | 8.717 |
| | CASP15 | 0.270 | 0.273 | 1.530 | 0.712 | 24.198 |
| | CASP16 | 0.217 | 0.218 | 1.686 | 0.343 | 41.482 |
| RiNALMo | RNA-Puzzles | 0.393 | 0.541 | 1.477 | 0.651 | 8.820 |
| | CASP15 | 0.240 | 0.231 | 1.506 | 0.732 | 26.210 |
| | CASP16 | 0.263 | 0.261 | 1.620 | 0.551 | 32.541 |
| trRosettaRNA | RNA-Puzzles | 0.460 | 0.602 | 1.424 | 0.655 | 7.264 |
| | CASP15 | 0.266 | 0.273 | 1.502 | 0.719 | 25.950 |
| | CASP16 | 0.284 | 0.284 | 1.543 | 0.575 | 30.328 |
| AlphaFold3 | RNA-Puzzles | 0.468 | 0.585 | 1.073 | 0.778 | 7.894 |
| | CASP15 | 0.374 | 0.345 | 1.035 | 0.936 | 18.979 |
| | CASP16 | 0.421 | 0.380 | 1.222 | 0.718 | 19.171 |

**Supplementary Table 4**: Entanglement analysis of tertiary structures predicted by structRFM for RNA-Puzzles (20 RNAs), CASP15 (12 RNAs) and CASP16 (13 RNAs). Entries are total counts across all predicted targets in each dataset for the nine entanglement types [2]: D&D, D&L, L&L, D(D), D(L), D(S), L(D), L(L) and L(S).

| Dataset | D&D | D&L | L&L | D(D) | D(L) | D(S) | L(D) | L(L) | L(S) |
|---|---|---|---|---|---|---|---|---|---|
| RNA-Puzzles | 10 | 3 | 4 | 5 | 14 | 0 | 3 | 0 | 1 |
| CASP15 | 28 | 7 | 1 | 16 | 15 | 0 | 8 | 0 | 0 |
| CASP16 | 3 | 2 | 1 | 6 | 3 | 1 | 1 | 0 | 0 |

**Supplementary Table 5**: Stereochemical-quality analysis of tertiary structures predicted by structRFM for RNA-Puzzles (20 RNAs), CASP15 (12 RNAs) and CASP16 (13 RNAs). Entries are total counts across all predicted targets in each dataset for six error types [3]: bond angle, bond length, chirality, close, planar and polymer.

| Dataset | bond angle | bond length | chirality | close | planar | polymer |
|---|---|---|---|---|---|---|
| CASP15 | 322 | 61 | 0 | 52 | 95 | 0 |
| CASP16 | 246 | 58 | 0 | 25 | 74 | 0 |
| RNA-Puzzles | 364 | 3 | 0 | 3 | 52 | 0 |

**Supplementary Table 6**: IRES-identification performance of fine-tuned structRFM and the comparison methods on the IRES test set (1,164 RNAs).

| Method | AUROC | F1 | Accuracy | Recall | Precision | MCC |
|---|---|---|---|---|---|---|
| structRFM | 0.713 | 0.701 | 0.650 | 0.821 | 0.612 | 0.320 |
| structRFM$^-$ | 0.684 | 0.678 | 0.650 | 0.737 | 0.628 | 0.305 |
| RiNALMo | 0.667 | 0.500 | 0.500 | 0.500 | 0.500 | 0.000 |
| IRESfinder | 0.449 | 0.475 | 0.475 | 0.474 | 0.475 | -0.050 |
| DeepIRES | 0.584 | 0.284 | 0.507 | 0.196 | 0.518 | 0.018 |

**Supplementary Table 7**: ncRNA-classification performance of fine-tuned structRFM and the comparison methods on nRC (2,600 RNAs). Results for the baseline methods are taken from RNAErnie [1].

| Method | Accuracy | Recall | Precision | F1 | MCC | AUROC |
|---|---|---|---|---|---|---|
| structRFM | 0.946 | 0.979 | 0.979 | 0.979 | 0.977 | 0.999 |
| structRFM$^{-}$ | 0.946 | 0.946 | 0.948 | 0.946 | 0.942 | 0.997 |
| RiNALMo | 0.889 | 0.889 | 0.912 | 0.891 | 0.882 | 0.995 |
| RNA-FM | 0.972 | 0.972 | 0.972 | 0.972 | 0.969 | 0.999 |
| RNA-MSM | 0.933 | 0.933 | 0.935 | 0.934 | 0.928 | 0.995 |
| RNABERT | 0.747 | 0.747 | 0.757 | 0.750 | 0.727 | 0.951 |
| RNAcon | 0.374 | 0.373 | 0.450 | 0.351 | 0.334 | - |
| nRC | 0.696 | 0.689 | 0.688 | 0.688 | 0.663 | - |
| ncRFP | 0.797 | 0.788 | 0.790 | 0.788 | 0.771 | - |
| RNAGCN | 0.857 | 0.861 | 0.988 | 0.856 | 0.846 | - |
| ncRDeep | 0.880 | 0.884 | 0.891 | 0.886 | 0.880 | - |
| ncRDense | 0.9510 | 0.951 | 0.953 | 0.951 | 0.947 | - |
| RNAErnie | 0.964 | 0.964 | 0.964 | 0.964 | 0.961 | - |

**Supplementary Table 8**: INF performance of structRFM and secondary-structure annotation baselines on ArchiveII600 [4] (3,911 RNAs) and bpRNA-TS0 [5] (1,305 RNAs).

| Method | ArchiveII600 | bpRNA-TS0 |
|---|---|---|
| ViennaRNA RNAfold | 0.579 | 0.522 |
| CONTRAfold | 0.557 | 0.597 |
| BPfold | 0.823 | 0.670 |
| structRFM-BPfold | 0.873 | 0.641 |
| structRFM-MUSES | 0.890 | 0.669 |

**Supplementary Table 9: Comparison of structRFM models pre-trained with BPfold- or MUSES-derived structure annotations across zero-shot, structure-prediction and function-prediction tasks.**

| **Task** | **Dataset** | **Number** | **Metric** | **Method** | **Mean** |
|---|---|---|---|---|---|
| Zero-shot homology classification | Rfam | 24,523 | OR ↓ | BPfold | **0.097** |
| | | | | MUSES | 0.099 |
| | ArchiveII600 | 3,911 | OR ↓ | BPfold | 0.483 |
| | | | | MUSES | **0.454** |
| Zero-shot structure prediction | PDB | 116 | INF ↑ | BPfold | 0.44 |
| | | | | MUSES | **0.46** |
| Secondary structure prediction | ArchiveII600 | 3,911 | INF ↑ | BPfold | 0.873 |
| | | | | MUSES | **0.890** |
| | bpRNA-TS0 | 1,305 | INF ↑ | BPfold | 0.641 |
| | | | | MUSES | **0.669** |
| Tertiary structure prediction | CASP15-natural | 8 | RMSD (Å) ↓ | BPfold | 14.8 |
| | | | | MUSES | **9.97** |
| | RNA-Puzzles | 20 | RMSD (Å) ↓ | BPfold | 6.82 |
| | | | | MUSES | **5.32** |
| | CASP15-natural | 8 | INF ↑ | BPfold | 0.66 |
| | | | | MUSES | **0.84** |
| | RNA-Puzzles | 20 | INF ↑ | BPfold | 0.65 |
| | | | | MUSES | **0.72** |
| Splice site prediction | SpliceAI | 16,505 | Top-k acc ↑ | BPfold | 0.42 |
| | | | | MUSES | 0.42 |
| IRES identification | IRES | 1,164 | F1 ↑ | BPfold | **0.708** |
| | | | | MUSES | 0.701 |
| ncRNA classification | nRC | 2,600 | F1 ↑ | BPfold | 0.973 |
| | | | | MUSES | **0.979** |

**Supplementary Table 10: Comparison of structRFM and structRFM$^-$ across zero-shot, structure-prediction and function-prediction tasks. Relative improvement is calculated as $|1 - \mathrm{Metric}_{\mathrm{structRFM}}/\mathrm{Metric}_{\mathrm{structRFM}^-}|$.**

| Task | Dataset | Number | Metric | structRFM$^-$ | structRFM | Improvement |
|---|---|---|---|---|---|---|
| Zero-shot | Rfam | 24,523 | OR ↓ | 0.115 | **0.099** | 13.9% |
| homology classification | ArchiveII600 | 3,911 | OR ↓ | 0.506 | **0.454** | 10.3% |
| Secondary structure | ArchiveII600 | 3,911 | INF ↑ | 0.852 | **0.890** | 4.46% |
| prediction | bpRNA-TS0 | 1,305 | INF ↑ | 0.630 | **0.669** | 6.19% |
| Tertiary structure | CASP15-natural | 8 | $\epsilon$RMSD ↓ | 1.48 | **1.26** | 14.9% |
| prediction | CASP16 | 13 | $\epsilon$RMSD ↓ | 1.69 | **1.49** | 11.8% |
| | RNA-Puzzles | 20 | $\epsilon$RMSD ↓ | 1.48 | **1.26** | 14.0% |
| | CASP15-natural | 8 | TM-score ↑ | 0.30 | **0.45** | 50.0% |
| | CASP16 | 13 | TM-score ↑ | 0.22 | **0.33** | 50.0% |
| | RNA-Puzzles | 20 | TM-score ↑ | 0.34 | **0.58** | 70.6% |
| | CASP15-natural | 8 | INF ↑ | 0.73 | **0.84** | 15.1% |
| | CASP16 | 13 | INF ↑ | 0.34 | **0.60** | 76.5% |
| | RNA-Puzzles | 20 | INF ↑ | 0.60 | **0.72** | 20.0% |
| Splice site prediction | SpliceAI | 16,505 | Top-k acc ↑ | 0.39 | **0.42** | 7.69% |
| IRES identification | IRES | 1,164 | F1 ↑ | 0.678 | **0.701** | 3.39% |
| ncRNA classification | nRC | 2,600 | F1 ↑ | 0.946 | **0.979** | 3.45% |

### Supplementary References


[1] Wang, N. *et al.* Multi-purpose rna language modelling with motif-aware pre-training and type-guided fine-tuning. *Nature Machine Intelligence* **6**, 548–557 (2024).

[2] Popenda, M. *et al.* Entanglements of structure elements revealed in rna 3d models. *Nucleic acids research* **49**, 9625–9632 (2021).

[3] Carrascoza, F., Antczak, M., Miao, Z., Westhof, E. & Szachniuk, M. Evaluation of the stereochemical quality of predicted rna 3d models in the rna-puzzles submissions. *Rna* **28**, 250–262 (2022).

[4] Sloma, M. F. & Mathews, D. H. Exact calculation of loop formation probability identifies folding motifs in rna secondary structures. *RNA* **22**, 1808–1818 (2016).

[5] Danaee, P. *et al.* bprna: large-scale automated annotation and analysis of rna secondary structure. *Nucleic acids research* **46**, 5381–5394 (2018).
